# Genomic prediction with whole-genome sequence data in intensely selected pig lines

**DOI:** 10.1101/2022.02.02.478838

**Authors:** Roger Ros-Freixedes, Martin Johnsson, Andrew Whalen, Ching-Yi Chen, Bruno D Valente, William O Herring, Gregor Gorjanc, John M Hickey

## Abstract

**Background:** Early simulations indicated that whole-genome sequence data (WGS) could improve genomic prediction accuracy and its persistence across generations and breeds. However, empirical results have been ambiguous so far. Large data sets that capture most of the genome diversity in a population must be assembled so that allele substitution effects are estimated with high accuracy. The objectives of this study were to use a large pig dataset to assess the benefits of using WGS for genomic prediction compared to using commercial marker arrays, to identify scenarios in which WGS provides the largest advantage, and to identify potential pitfalls for its effective implementation.

**Methods:** We sequenced 6,931 individuals from seven commercial pig lines with different numerical size. Genotypes of 32.8 million variants were imputed for 396,100 individuals (17,224 to 104,661 per line). We used BayesR to perform genomic prediction for eight complex traits. Genomic predictions were performed using either data from a marker array or variants preselected from WGS based on association tests.

**Results:** The prediction accuracy with each set of preselected WGS variants was not robust across traits and lines and the improvements in prediction accuracy that we achieved so far with WGS compared to marker arrays were generally small. The most favourable results for WGS were obtained when the largest training sets were available and used to preselect variants with statistically significant associations to the trait for augmenting the established marker array. With this method and training sets of around 80k individuals, average improvements of genomic prediction accuracy of 0.025 were observed in within-line scenarios.

**Conclusions:** Our results showed that WGS has a small potential to improve genomic prediction accuracy compared to marker arrays in intensely selected pig lines in some settings. Thus, although we expect that more robust improvements could be attained with a combination of larger training sets and optimised pipelines, the use of WGS in the current implementations of genomic prediction should be carefully evaluated on a case-by-case basis against the cost of generating WGS at a large scale.

## Introduction

Whole-genome sequence data (WGS) has the potential to empower the identification of causal variants that underlie quantitative traits or diseases [1–4], increase the precision and scope of population genetic studies [5,6], and enhance livestock breeding. Genomic prediction has been successfully implemented in the main livestock species and it has increased the rate of genetic gain [7]. Genomic prediction has provided many benefits such as greater accuracies of genetic evaluations in livestock populations, such as cattle and pig, and the reduction of the generational interval, most notably in dairy cattle. Since its early implementations, genomic prediction is typically performed using marker arrays that capture the effects of the (usually unknown) causal variants via linkage and linkage disequilibrium [8,9]. In contrast, WGS are assumed to contain the causal variants. For this reason, it was hypothesized that WGS could further improve genomic prediction accuracy and its persistence across generations and breeds. Early simulations indicated that causal mutations from WGS could increase prediction accuracy. One simulation study indicated that the magnitude of prediction accuracy improvement relative to dense marker arrays ranged from 2.5 to 3.7%, with a persistence of over 10 generations [10]. Another study reported improvements in prediction accuracy of up to 30% if causal variants with low minor allele frequency could be captured by the WGS [11]. However, benefits could be low in typical livestock populations due to small effective population sizes and recent directional selection [12].

During the last few years, there have been several attempts at improving the accuracy of genomic prediction with WGS in the main livestock species. Empirical results have been ambiguous so far. When predicting genomic breeding values within a population, some studies found no relevant improvement in genomic prediction accuracy with WGS compared to marker arrays [13–16]. Other studies found small, and often unstable, improvements (e.g., from 1 to 5% or no improvement depending on prediction method [17–19], or trait-dependent results [19,20]). With genomic prediction across populations, the identification of causal variants from WGS can improve prediction accuracy [21–24], especially for numerically small populations or for populations that are not represented in the training [21,23–27].

One of the most successful strategies to exploit WGS consists in augmenting available marker arrays with preselected variants from WGS based on their association with the trait of interest [28–31]. In some cases, this strategy improved genomic prediction accuracy by up to 9% [30] and 11% [31], but this strategy did not improve prediction accuracy in other within-line settings [15]. Nevertheless, these examples indicate how identifying causal variants could enhance genomic prediction with WGS. Whole-genome sequence data has already been applied in genome-wide association studies (GWAS) to identify variants associated to a variety of traits in livestock [2,32–34], including pigs [35,36]. However, the fine-mapping of causal variants remains challenging due to the pervasive long-range linkage disequilibrium across extremely dense variation [37].

The estimation of allele substitution effects with high accuracy and, ideally, the identification of causal variants amongst millions of other variants are important for the usefulness of WGS in research and breeding. This requires large data sets able to capture most of the genome diversity in a population. Low-cost sequencing strategies have been developed, which typically involve sequencing a subset of the individuals in a population at low coverage and then imputing WGS for the remaining individuals. However, despite this, the cost of generating accurate WGS at such a large scale, as well as the large computational requirements for the analyses of such datasets, have limited the population sizes or number of populations tested in some of the previous studies. This hinders the interpretation of results across studies, which are very diverse in population structures, sequencing strategies and prediction methodologies used. The largest studies in livestock on the use of WGS for genomic prediction to date have been performed in cattle, for which a large multi-breed reference panel is available from the 1000 Bull Genomes Project [2,17,32]. This reference panel has enabled the imputation of WGS in many cattle populations. The lack of such reference panels hampers the potential of WGS in other species, such as pigs [35].

We have previously described our approach to impute WGS in large pedigreed populations without external reference panels [38]. Following that strategy, we generated WGS for 396,100 pigs from seven intensely selected lines with diverse genetic backgrounds and numerical size. The objectives of this study were to use this large pig dataset to assess the benefits of using WGS for genomic prediction compared to using commercial marker arrays, to identify scenarios in which WGS provides the largest advantage, and to identify potential pitfalls for its effective implementation.

## Materials and Methods

### Populations and sequencing strategy

We re-sequenced the whole genome of 6,931 individuals from seven commercial pig lines (Genus PIC, Hendersonville, TN) with a total coverage of approximately 27,243x. Breeds of origin of the nine lines included Large White, Landrace, Pietrain, Hampshire, Duroc, and synthetic lines. Sequencing effort in each of the seven lines was proportional to population size. The number of pigs that were available in the pedigree of each line and the number of sequenced pigs, by coverage, is summarized in Table 1. Approximately 1.5% (0.9 to 2.1% in each line) of the pigs in each line were sequenced. Most pigs were sequenced at low coverage, with target coverage of 1 or 2x, but a subset of pigs was sequenced at a higher coverage of 5, 15, or 30x. Thus, the average individual coverage was 3.9x, but the median coverage was 1.5x. Most of the sequenced pigs were born during the 2008–2016 period. The population structure across the seven lines was assessed with a principal component analysis using the sequenced pigs and is shown in Additional file 1.

**Table 1.** Number of sequenced pigs and pigs with imputed data.

| Line | Individuals<br>sequenced | Individuals sequenced by<br>coverage |  |  |  | Individuals used in the analyses |  |  |  |
| --- | --- | --- | --- | --- | --- | --- | --- | --- | --- |
|  |  | 1x | 2x | 5x | 15–30x | Pedigree | LD | HD | Imputed |
| A | 1,856 | 1,044 | 649 | 73 | 90 | 122,753 | 39,485 | 66,763 | 104,661 |
| B | 1,366 | 685 | 545 | 44 | 92 | 88,964 | 39,110 | 38,763 | 76,230 |
| C | 1,491 | 628 | 728 | 54 | 81 | 84,420 | 35,343 | 34,358 | 66,608 |
| D | 731 | 362 | 311 | 16 | 42 | 79,981 | 16,650 | 54,297 | 60,474 |
| E | 760 | 394 | 274 | 27 | 65 | 50,797 | 22,768 | 20,685 | 41,573 |
| F | 381 | 193 | 137 | 16 | 35 | 35,309 | 11,747 | 17,758 | 29,330 |
| G | 445 | 217 | 176 | 15 | 37 | 21,129 | 11,472 | 6,661 | 17,224 |

*Pedigree* number of individuals included in the pedigree used for imputation, *LD*
number of individuals genotyped with the low-density marker array, *HD* number of
individuals genotyped with high-density marker arrays, *Imputed* number of
individuals with imputed genotypes that remain after filtering out individuals with
low predicted imputation accuracy.

The sequenced pigs and their coverage were selected following a three-part sequencing strategy developed to represent the haplotype diversity in each line. First (1), sires and dams with the highest number of genotyped progeny were sequenced at 2x and 1x, respectively. Sires were sequenced at a greater coverage because they contributed with more progeny than dams. Then (2), the individuals with the greatest genetic footprint on the population (i.e., those that carry more of the most common haplotypes) and their immediate ancestors were sequenced at a coverage between 1x and 30x (AlphaSeqOpt part 1; [39]). The sequencing coverage was allocated with an algorithm that maximises the expected phasing accuracy of the common haplotypes from the accumulated family information. Finally (3), pigs that carried haplotypes with low accumulated coverage (below 10x) were sequenced at 1x (AlphaSeqOpt part 2; [40]). Sets (2) and (3) were based on haplotypes inferred from marker array genotypes (GGP-Porcine HD BeadChip; GeneSeek, Lincoln, NE), which were phased with AlphaPhase [41] and imputed with AlphaImpute [42].

Most sequenced pigs and their relatives were also genotyped with marker arrays either at low density (15k markers) using the GGP-Porcine LD BeadChip (GeneSeek) or at high density (50k or 80k markers) using different versions of the GGP-Porcine HD BeadChip (GeneSeek). In our study we only used markers included in the 50k array, which is the latest version of the high-density array. Markers in the 15k array were nested within the 50k array and markers from the 80k array that were not included in the 50k array were discarded. The number of pigs genotyped at each density is summarized in Table 1. Quality control of the marker array data was based on the individuals genotyped at high density. Markers with minor allele frequency below 0.01, call rate below 0.80, or a significant deviation from the Hardy-Weinberg equilibrium were removed. After quality control, 38,634 to 43,966 markers remained in each line.

### Sequencing and data processing

Tissue samples were collected from ear punches or tail clippings. Genomic DNA was extracted using Qiagen DNeasy 96 Blood & Tissue kits (Qiagen Ltd., Mississauga, ON, Canada). Paired-end library preparation was conducted using the TruSeq DNA PCR-free protocol (Illumina, San Diego, CA). Libraries for resequencing at low coverage (1 to 5x) were produced with an average insert size of 350 bp and sequenced on a HiSeq 4000 instrument (Illumina). Libraries for resequencing at high coverage (15 or 30x) were produced with an average insert size of 550 bp and sequenced on a HiSeq X instrument (Illumina). All libraries were sequenced at Edinburgh Genomics (Edinburgh Genomics, University of Edinburgh, Edinburgh, UK).

DNA sequence reads were pre-processed using Trimmomatic [43] to remove adapter sequences from the reads. The reads were then aligned to the reference genome *Sscrofa11.1* (GenBank accession: GCA_000003025.6) using the BWA-MEM algorithm [44]. Duplicates were marked with Picard (http://broadinstitute.github.io/picard). Single nucleotide polymorphisms (SNPs) and short insertions and deletions (indels) were identified with the variant caller GATK HaplotypeCaller (GATK 3.8.0) [45,46] using default settings. Variant discovery with GATK HaplotypeCaller was performed separately for each individual and then a joint variant set for all the individuals in each population was obtained by extracting the variant positions from all the individuals.

We extracted the read counts supporting each allele directly from the aligned reads stored in the BAM files using a pile-up function to avoid biases towards the reference allele introduced by GATK when applied on low-coverage WGS [47]. That pipeline uses pysam (version 0.13.0; https://github.com/pysam-developers/pysam), which is a wrapper around htslib and the samtools package [48]. We extracted the read counts for all biallelic variants, after filtering out variants observed in less than three sequenced individuals and variants in potential repetitive regions (defined as variants that had mean depth values 3 times greater than the average realized coverage) with VCFtools [49]. This pipeline delivered a total of 55.6 million SNP (19.6 to 31.1 million within each line) and 10.2 million indels (4.1 to 5.6 million within each line). A more complete description of the variation across the lines is provided in [50].

### Genotype imputation

Genotypes were jointly called, phased and imputed for a total of 483,353 pedigree-related individuals using the ‘hybrid peeling’ method implemented in AlphaPeel [51,52]. This method used all the available marker array and WGS. Imputation was performed separately for each line using complete multi-generational pedigrees, with 21,129 to 122,753 individuals per line (Table 1). We have previously published reports on the accuracy of imputation in the same populations using this method [38]. The estimated average individual-wise dosage correlation was 0.94 (median: 0.97). Individuals with low predicted imputation accuracy were removed before further analyses. An individual was predicted to have low imputation accuracy if itself or all of its grandparents were not genotyped with a marker array or if it had a low degree of connectedness to the rest of the population (defined as the sum of coefficients of pedigree-based relationship between the individual and the rest of individuals). These criteria were based on the analysis of imputation accuracy in simulated and empirical data [38]. A total of 396,100 individuals remained, with 17,224 and 104,661 individuals per line (Table 1). The expected average individual-wise dosage correlation of the remaining individuals was 0.97 (median: 0.98) according to our previous estimates. We also excluded from the analyses variants with a minor allele frequency lower than 0.023, because their estimated variant-wise dosage correlations was lower than 0.90 [38]. After imputation, 32.8 million variants (14.5 to 19.9 million within each line) remained for downstream analyses, out of which 9.9 million segregated across all seven lines.

### Traits

We analysed data of eight complex traits that are commonly included in selection objectives of pig breeding programmes: average daily gain (ADG, g), backfat thickness (BFT, mm), loin depth (LD, mm), average daily feed intake (ADFI, kg), feed conversion ratio (FCR), total number of piglets born (TNB), litter weight at weaning (LWW, kg), and return to oestrus 7 days after weaning (RET, binary trait). Most pigs with records were born during the 2008–2020 period. Breeding values were estimated by line with a linear mixed model that included polygenic and non-genetic (as relevant for each trait) effects. Deregressed breeding values (dEBV) were obtained following the method of VanRaden et al. [53]. Only individuals in which the trait was directly measured were retained for further analyses. The number of records for each trait used in the analyses of each line is detailed in Table 2.

**Table 2.** Number of phenotype records per trait and line.

| **Trait** | **Set** | **A** | **B** | **C** | **D** | **E** | **F** | **G** |
| --- | --- | --- | --- | --- | --- | --- | --- | --- |
| ADG | Training | 77,811 | 54,709 | 48,219 | 45,693 | 31,918 | 24,046 | 13,479 |
|  | Test | 9,435 | 8,387 | 6,977 | 4,789 | 3,019 | 1,808 | 1,572 |
| BFT | Training | 70,529 | 52,910 | 47,512 | 42,636 | 31,127 | 21,892 | 13,300 |
|  | Test | 8,560 | 7,957 | 6,747 | 4,301 | 2,936 | 1,602 | 1,568 |
| LD | Training | 75,117 | 54,537 | 48,054 | 43,517 | 31,987 | 24,154 | 13,303 |
|  | Test | 9,021 | 8,415 | 6,995 | 4,411 | 3,024 | 1,807 | 1,570 |
| ADFI | Training | 20,535 | 8,866 | 8,235 | 11,573 | 11,930 | 4,000 | 4,457 |
|  | Test | 1,358 | 638 | 802 | 641 | 478 | 97* | 364 |
| FCR | Training | 19,805 | 8,572 | 7,857 | 11,378 | 11,804 | 3,900 | 4,364 |
|  | Test | 1,328 | 624 | 775 | 624 | 477 | 97* | 360 |
| TNB | Training | 12,250 | 9,315 | 8,438 | 7,700 | 5,834 | - | 2,865 |
|  | Test | 254 | 428 | 400 | 23* | 125* | - | 98* |
| LWW | Training | - | 7,884 | 6,251 | - | - | - | 2,505 |
|  | Test | - | 246 | 220 | - | - | - | 47* |
| RET | Training | - | 5,928 | 5,496 | - | - | - | 1,481 |
|  | Test | - | 332 | 282 | - | - | - | 70* |

*ADG* average daily gain, *BFT* backfat thickness, *LD* loin depth, *ADFI* average daily
feed intake, *FCR* feed conversion ratio, *TNB* total number of piglets born, *LWW* litter
weight at weaning, *RET* return to oestrus 7 days after weaning.
\*Included in multi-line scenarios, but excluded in within-line scenarios because of the
limited size of the testing set.

### Training and testing sets

We split the individuals in each line into training and testing sets. The testing sets were defined as individuals from full-sib families from the last generation of the pedigree (i.e., individuals that did not have any progeny of their own). Only families with a minimum of 5 full-sibs were considered. The training set was defined as all those individuals that had a pedigree coefficient of relationship lower than 0.5 with any individual in the testing set. This design was chosen to mimic a realistic situation in which breeding programmes evaluate selection candidates available in a selection nucleus at any given time.

To assess the effect of the size of the training set on prediction accuracy, we created training sets with a reduced number of phenotype records for the three largest lines and the three traits with the largest number of records. We did this by removing the oldest animals in a way that approximately the most recent 10, 20, or 35 to 45 thousand phenotype records remained in each of the reduced training sets.

Due to the computational requirements of the analyses, we could not perform repetitions for every analysis. However, we estimated variability of the results across repetitions in the largest, an intermediate, and the smallest lines for two traits with a large and small number of phenotype records. For doing this, we randomly split the test sets into five subsets, with each full-sib family represented exclusively in one of the subsets. Training sets for each repetition were defined as for the general case.

### Genome-wide association study

To provide an association-based criterion to preselect variants for genomic prediction, we performed a GWAS for each trait and line. This step included only the individuals in the training set. We fitted a univariate linear mixed model that accounted for the genomic relationships as:

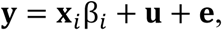

where **y** is the vector of dEBV, **x***_i_* is the vector of genotypes for the *i*th variant coded as 0 and 2 if homozygous for either allele or 1 if heterozygous, β*_i_* is the allele substitution effect of the *i*th variant on the trait, 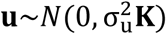 is the vector of polygenic effects with the covariance matrix equal to the product of the polygenic additive genetic variance 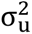 and a genomic relationship matrix **K**, and **e** is a vector of uncorrelated residuals. Due to computational limitations, the genomic relationship matrix **K** was calculated using only imputed genotypes in the marker array. We used the FastLMM software [54,55] to fit the model.

### Within-line genomic prediction

To test whether variants from the WGS could provide greater genomic prediction accuracy than the marker array, we tested genomic prediction using variants from the marker array, from the WGS, or combining them. The marker array data (also referred to as ‘Chip’) was set as the benchmark for prediction accuracy. It contained all ∼40k variants in the marker array. For WGS, we preselected sets of variants because currently available methods for genomic prediction are not yet capable of handling datasets as large as the complete WGS without exorbitant computational resources. We tested different alternative strategies for preselecting variants for the prediction model based on the GWAS results:

- *Top40k.* To mimic the number of variants in Chip, we preselected the variants with the lowest p-value (not necessarily below the significance threshold) in each of consecutive non-overlapping 55-kb windows along the genome. In addition, to test the impact of variant density on prediction accuracy, we preselected 10k, 25k, 75k, or 100k variants following the same criterion.
- *ChipPlusSign.* Variants preselected as in Top40k, but only significant variants (p≤10^-6^) were preselected and merged with those in Chip. When a 55-kb window contained more than one significant variant, only that with the lowest p-value was selected as a proxy to reduce the preselection of multiple significant variants tagging the same causal variant. When the most significant variant from WGS was already included in the marker array, the variant was considered only once and in the rare cases of genotype discordance, the genotype was replaced with the mean genotype value in that line. On average, 309 significant variants were identified per trait and line (range: 23 to 1083; Table 3) and merged with those in Chip.

**Table 3.** Number of significant variants from the whole-genome sequence data that were added to the marker array in ChipPlusSign.

| Trait | A | B | C | D | E | F | G | Multi-line |
| --- | --- | --- | --- | --- | --- | --- | --- | --- |
| ADG | 646 | 581 | 424 | 498 | 279 | 219 | 143 | 4731 |
| BFT | 1083 | 758 | 664 | 518 | 1030 | 218 | 237 | 6149 |
| LD | 633 | 579 | 458 | 518 | 222 | 215 | 43 | 7247 |
| ADFI | 145 | 224 | 169 | 23 | 183 | - | 119 | 767 |
| FCR | 198 | 224 | 162 | 95 | 56 | - | 134 | 1369 |
| TNB | 71 | 117 | 161 | - | - | - | - | 248 |
| LWW | - | 32 | 73 | - | - | - | - | 480 |
| RET | - | 184 | 31 | - | - | - | - | 60 |

*ADG* average daily gain, *BFT* backfat thickness, *LD* loin depth, *ADFI* average daily feed intake, *FCR* feed conversion ratio, *TNB* total number of piglets born, *LWW* litter weight at weaning, *RET* return to oestrus 7 days after weaning.

Genomic prediction was performed by fitting a univariate model with BayesR [56,57], which uses a mixture of normal distributions as the prior for variant effects, including one distribution that sets the variant effects to zero. The model was:

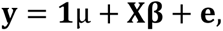

where **y** is the vector of dEBV, **1** is a vector of ones, μ is the general mean, **X** is a matrix of variant genotypes, **β** is a vector of variant effects, and **e** is a vector of uncorrelated residuals. The prior variance of the variant effects in **β** had four components with mean zero and variances 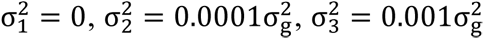 and 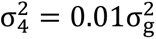, where 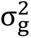 is the total genetic variance. We used a uniform and almost uninformative prior for the mixture distribution with the total genetic variance re-estimated in every iteration. We used a publicly available implementation of BayesR (https://github.com/syntheke/bayesR; accessed on 30 April 2021), with default settings. Prediction accuracy was calculated in the testing set as the correlation between the predicted genomic breeding values and the dEBV. Bias of the prediction accuracy was calculated as the regression coefficient of the dEBV on the predicted genomic breeding values. For ease of comparison between traits and lines, the difference between prediction accuracy of WGS and the marker array was calculated. The difference in prediction accuracy was analysed by fitting linear models with the size of the training set as a covariate and trait and line as fixed effects when appropriate.

### Multi-line genomic prediction

We considered multi-line scenarios in which the training set was formed by merging the training sets that had been defined for each line. All analyses were performed as for the within-line scenarios but with an additional effect of the line in the prediction model. In the multi-line scenarios, all variants from the marker array that passed quality control and were imputed for at least one line were included in the baseline (referred to as ‘ML-Chip’). For ease of computation, the strategies for preselection of variants from WGS were applied only to the subset of 9.9 million variants that had been called and imputed in all seven lines. Thus, we defined the variant sets ‘ML-Top40k’ and ‘ML-ChipPlusSign’ by preselecting variants following the same criteria as in within-line scenarios, but using a multi-line GWAS with an additional effect of the line. For ML-ChipPlusSign, 60 to 7247 significant variants were identified per trait (Table 3) and merged with those in ML-Chip. For comparison purposes, genomic prediction accuracy was calculated for the testing set of each individual line.

## Results

### Within-line genomic prediction accuracy

Whole-genome sequence data improved genomic prediction accuracy compared to marker array data in some scenarios, especially when there was a sufficiently large training set and if an appropriate set of variants was preselected. Figure 1 shows the prediction accuracy for the three traits and three lines with the largest training sets using the two different sets of WGS variants. Results for the rest of traits and lines, as well as results for the bias, are provided in Additional File 2. For BFT in line B, the two tested sets of variants from the WGS increased prediction accuracy by 0.054 (+9.8%), for Top40k, and by 0.043 (+7.7%), for ChipPlusSign. However, the performance of WGS was not robust and differed for each trait and line, and even across repetitions within trait and line (Additional File 3), often leading to no improvements of prediction accuracy or even reduced prediction accuracy relative to the marker array. For instance, Top40k reduced prediction accuracy by 0.020 (– 3.4%), for ADG in line C, and ChipPlusSign by 0.020 (–2.2%), for LD in line C. Using WGS reduced bias compared to the marker array in some, but not all, scenarios.

**Figure 1.**
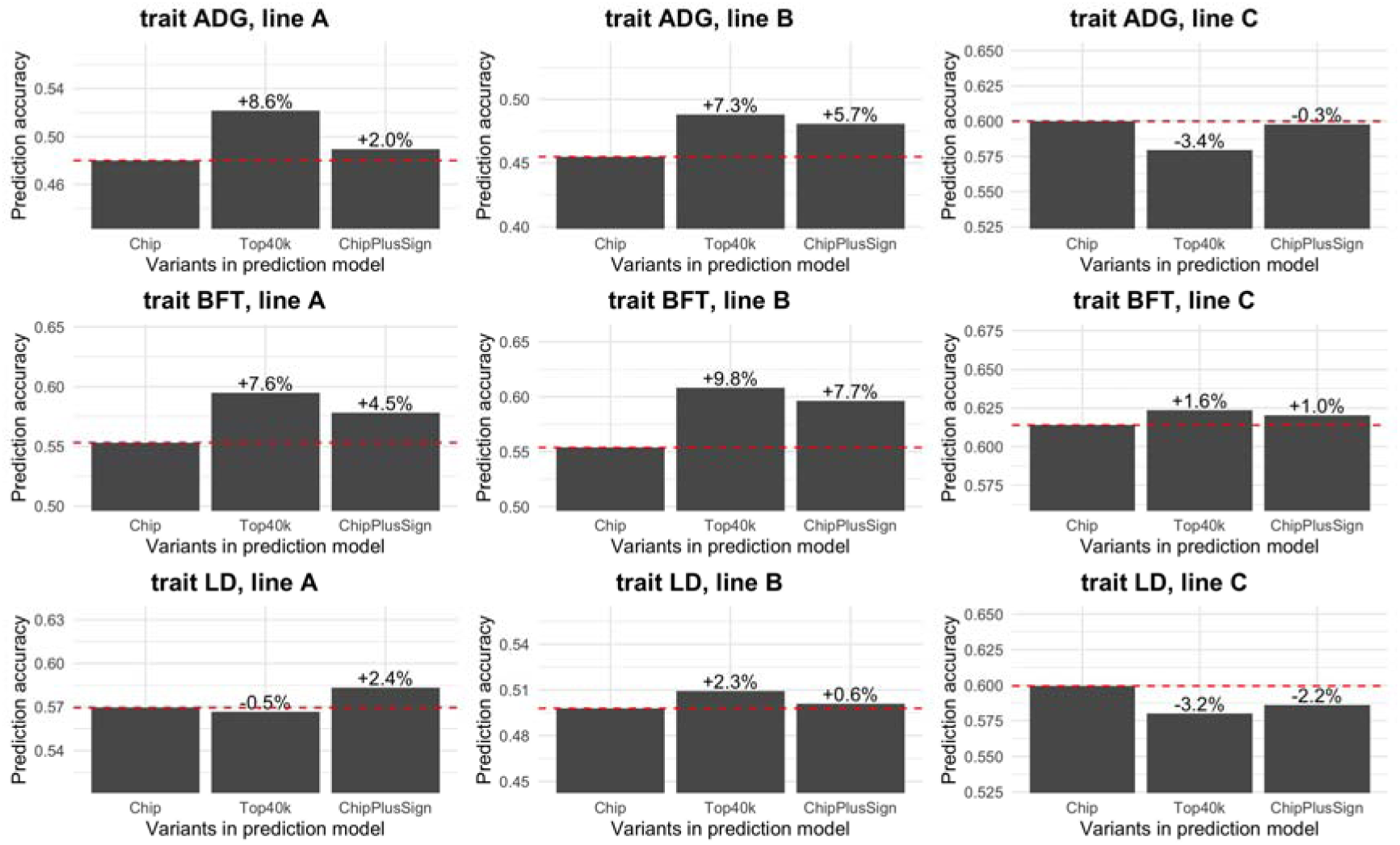
Genomic prediction accuracy for each set of variants for traits ADG, BFT, and LD in the three largest lines. Dashed line at value of marker array (Chip) as a reference. Values indicate relative difference to marker array (Chip).

There was a trend that the capacity of WGS variants to improve the genomic prediction accuracy compared to the marker arrays was larger for the traits and lines with larger training sets. Figures 2 and 3 show the difference in prediction accuracy of Top40k and ChipPlusSign compared to the marker array against the training set size. We observed large variability for the difference in prediction accuracy, especially when the training set was small. This variability was larger in Top40k than in ChipPlusSign, in a way that shrinkage of variation as the training set was larger was more noticeable in ChipPlusSign. Within trait and line, the variability across repetitions was also larger in Top40k than in ChipPlusSign (Additional File 3). Gains in prediction accuracy were low-to-moderate in the most favourable cases. In the most unfavourable cases we observed large losses in prediction accuracy for Top40k but more limited losses for ChipPlusSign with moderate training set sizes. The regression coefficient between the difference in prediction accuracy and the size of the training set was positive but had stronger statistical evidence for ChipPlusSign (b=0.5·10^-6^ individual^-1^; p=0.032; R^2^ for each trait between 0.06 and 0.75) than for Top40k (b=0.5·10^-6^ individual^-1^; p=0.24; R^2^ for each trait between 0.00 and 0.20), because of the apparent lower robustness with Top40k.

**Figure 2.**
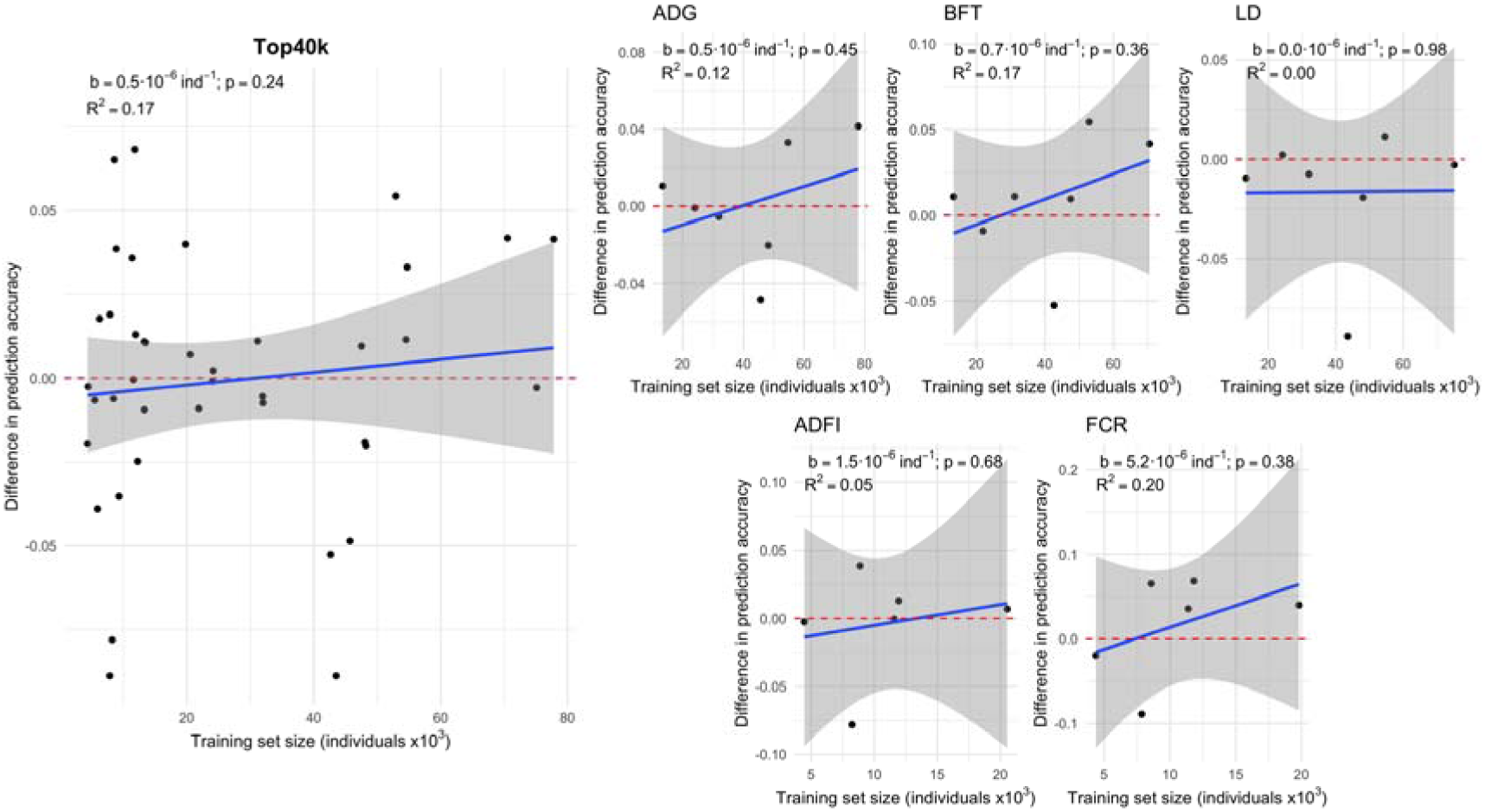
Genomic prediction accuracy with the Top40k variants for the complex traits. The difference between the Top40k and marker array is shown for all traits and lines (left) or by trait (right). Red dashed line at ‘no difference’. Regression coefficient (b) and p-value of training set size is provided, as well as the coefficient of determination (R^2^) of the model. The linear model for the joint analyses included the trait effect.

**Figure 3.**
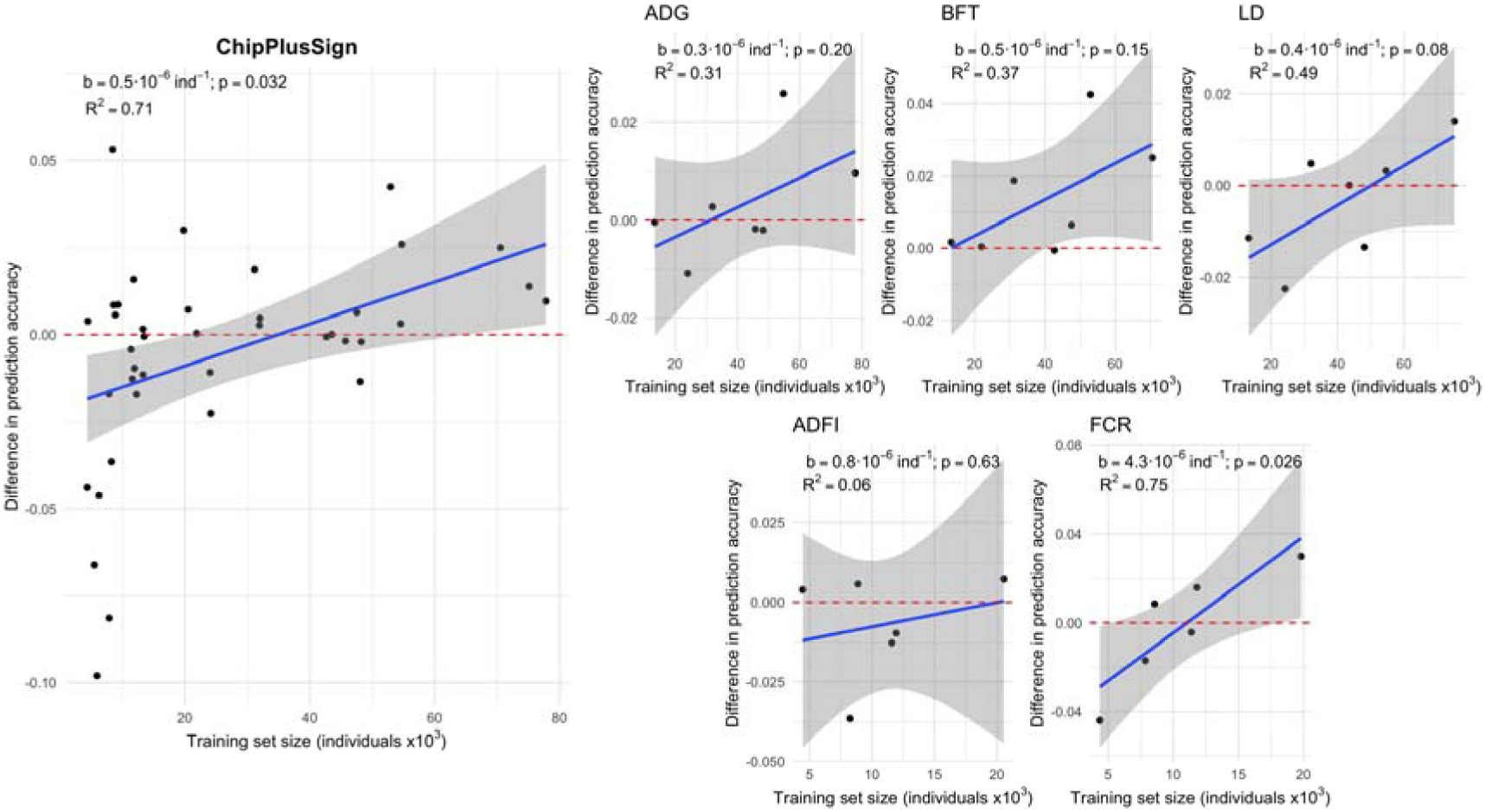
Genomic prediction accuracy with the ChipPlusSign variants for the complex traits. The difference between the ChipPlusSign and marker array is shown for all traits and lines (left) or by trait (right). Red dashed line at ‘no difference’. Regression coefficient (b) and p-value of training set size is provided, as well as the coefficient of determination (R^2^) of the model. The linear model for the joint analyses included the trait effect.

Results within trait and line (Figure 4) confirmed that the impact of WGS on genomic prediction accuracy depended on line, but also that in general WGS yielded higher prediction accuracy compared to the marker array when the training set was the largest. Under this setting, the regression coefficient between the difference in prediction accuracy and the size of training set was 0.6·10^-6^ individual^-1^ (p<0.001), for Top40k, and 0.3·10^-6^ individual^-1^ (p=0.017) for ChipPlusSign. This was at least partly driven by the lower number of significant associations that were detected with smaller training sets. With a training set of 20k individuals or less, 118 to 287 significant variants were added to the marker array; with a training set of 35k to 45k individuals, 288 to 709 significant variants; and with all available individuals in the training set, 424 to 1083 significant variants. Thus, if the marker array was augmented with the significant variants detected with all available individuals (ChipPlusSign*), then WGS yielded the same prediction accuracy than the marker array or higher in most scenarios even when the set for training the predictive equation was smaller.

**Figure 4.**
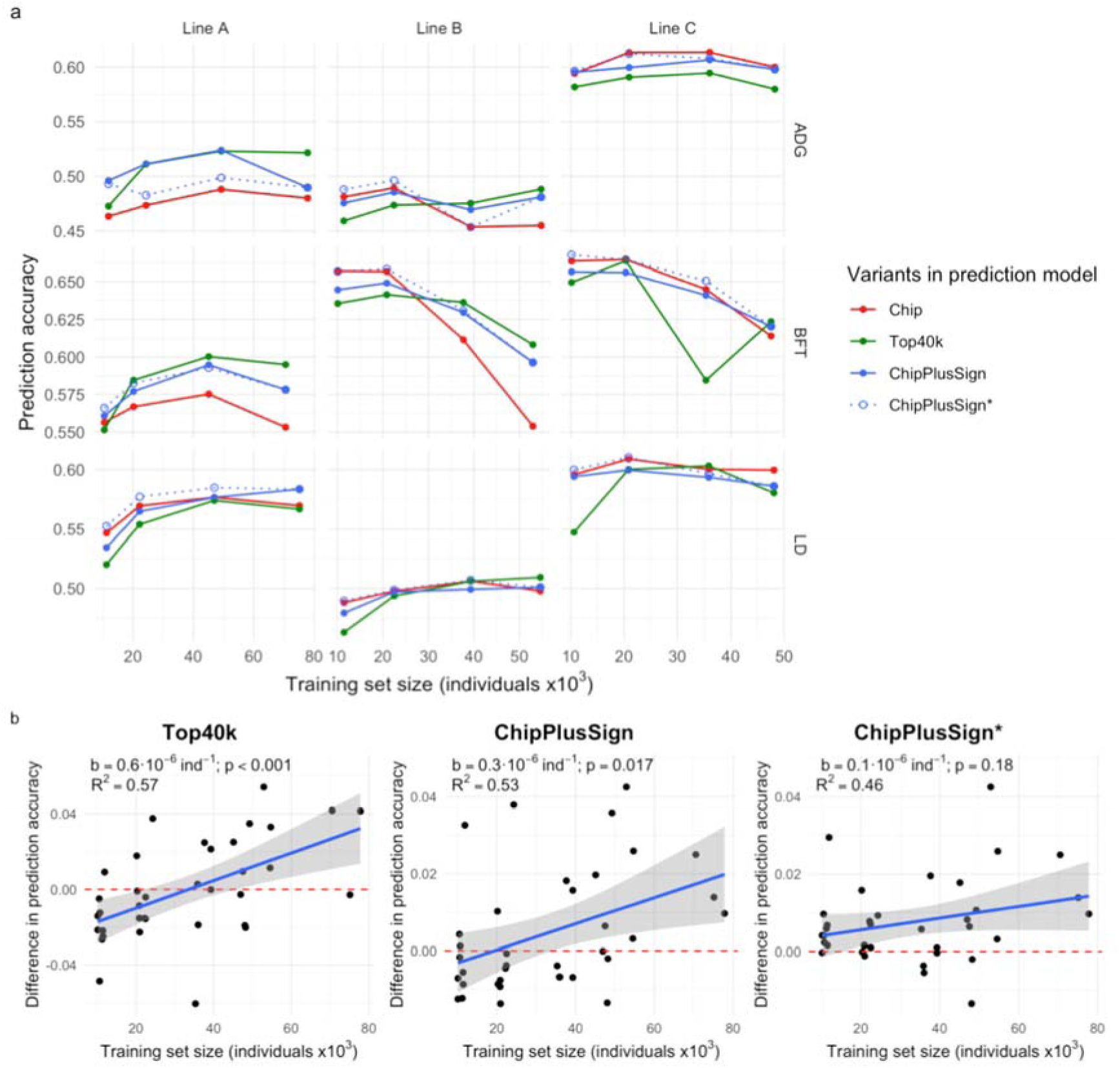
Effect of training set size on the genomic prediction accuracy for each set of variants for traits ADG, BFT, and LD in the three largest lines. **(a)** Genomic prediction accuracy with the marker array (Chip) or with preselected WGS data (Top40k, ChipPlusSign, and ChipPlusSign*). In ChipPlusSign* variants are preselected based on associations tested using the largest training set available. **(b)** The difference between the ChipPlusSign and Chip is shown for all traits and lines. Red dashed line at ‘no difference’. Regression coefficient (b) and p-value of training set size is provided, as well as the coefficient of determination (R^2^) of the model. The linear model for the joint analyses included the trait and line effects.

Results from simulated traits (Additional File 4) confirmed the trends observed for the empirical traits; for instance, the higher robustness of ChipPlusSign compared to Top40k. Results from the simulated traits also showed the impact of the genetic architecture of the traits on the success of WGS in improving genomic prediction accuracy. Traits with high heritability and low number of QTN were more likely to show larger improvements in prediction accuracy.

We observed diminishing returns when we increased the density of the variants used in prediction. Increasing the number of variants from the 40k in Top40k to 75k selected in the same way yielded small improvements in genomic prediction accuracy compared to Top40k, but increases up to 100k variants provided smaller or null additional gains (Additional File 5).

### Multi-line genomic prediction accuracy

The accuracy of genomic prediction trained across multi-line datasets was systematically lower than in the within-line datasets (Additional File 6). Nonetheless, when using multi-line training sets, the ML-ChipPlusSign variants in general increased genomic prediction accuracy relative to the marker array (ML-Chip; Figure 5). For the traits that accumulated the largest multi-line training sets (i.e., ADG, BFT, and LD), the improvements of prediction accuracy in each individual line seemed unrelated to the number of individuals that each line contributed to the multi-line training set. However, for the traits that accumulated smaller multi-line training sets (i.e., ADFI and FCR), ML-ChipPlusSign only improved prediction accuracy in the lines that contributed more individuals to the multi-line training set, and reduced prediction accuracy in the lines that contributed less individuals to the multi-line training set. Therefore, as happened in the within-line scenarios, the greatest improvements of prediction accuracy with WGS were achieved for the largest individual lines, although ML-ChipPlusSign in the multi-line scenarios also improved prediction accuracy compared to ML-Chip for some traits and lines for which no improvements were observed in the within-line scenarios, including numerically small lines (Figure 6). In contrast, results for ML-Top40k were not robust across traits (Additional File 7).

**Figure 5.**
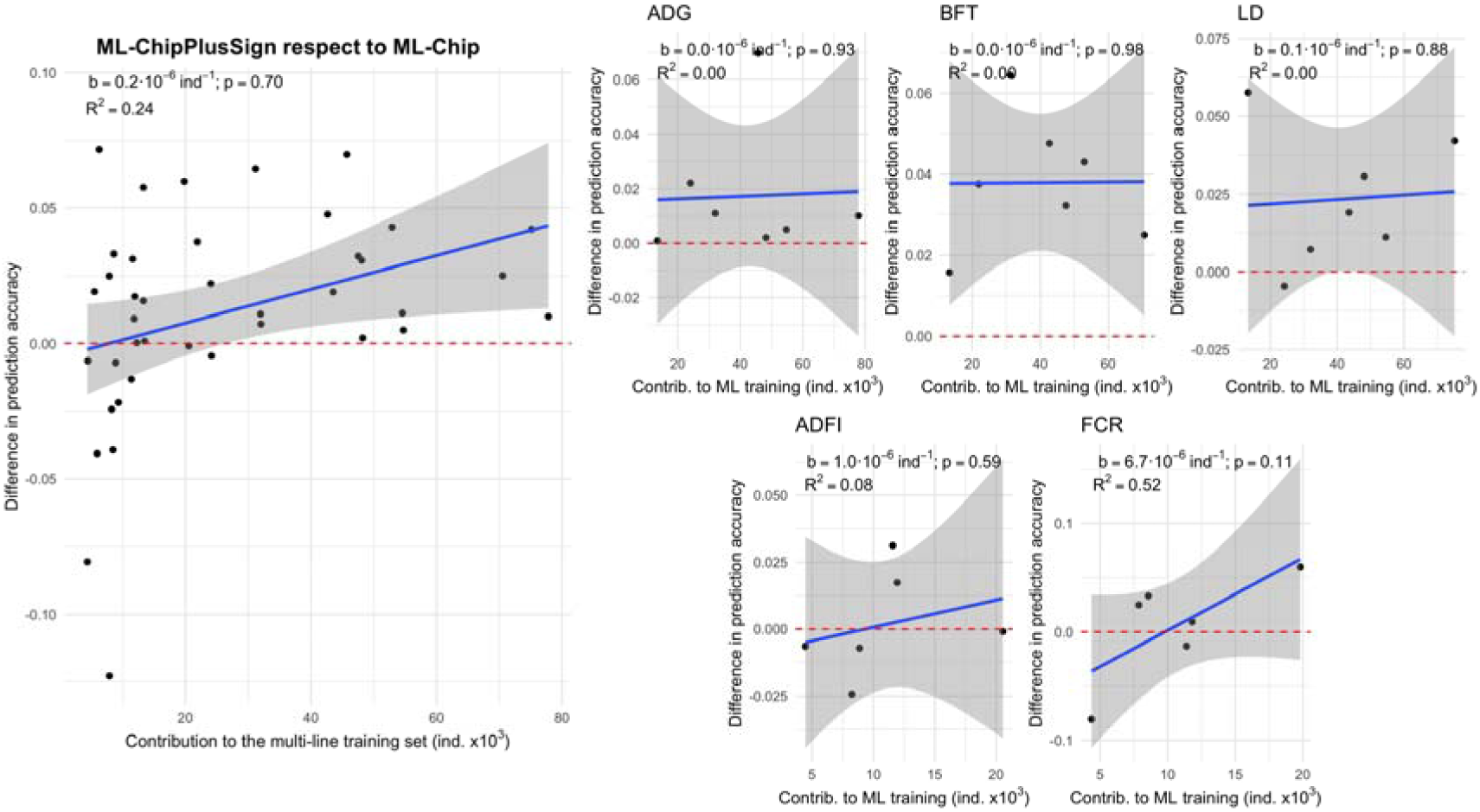
Genomic prediction accuracy with the ML-ChipPlusSign variants for the complex traits. The difference between ML-ChipPlusSign and marker array (ML-Chip) is shown for all traits and lines (left) or by trait (right). Red dashed line at ‘no difference’. Regression coefficient (b) and p-value of training set size is provided, as well as the coefficient of determination (R^2^) of the model. The linear model for the joint analyses included the trait effect.

**Figure 6.**
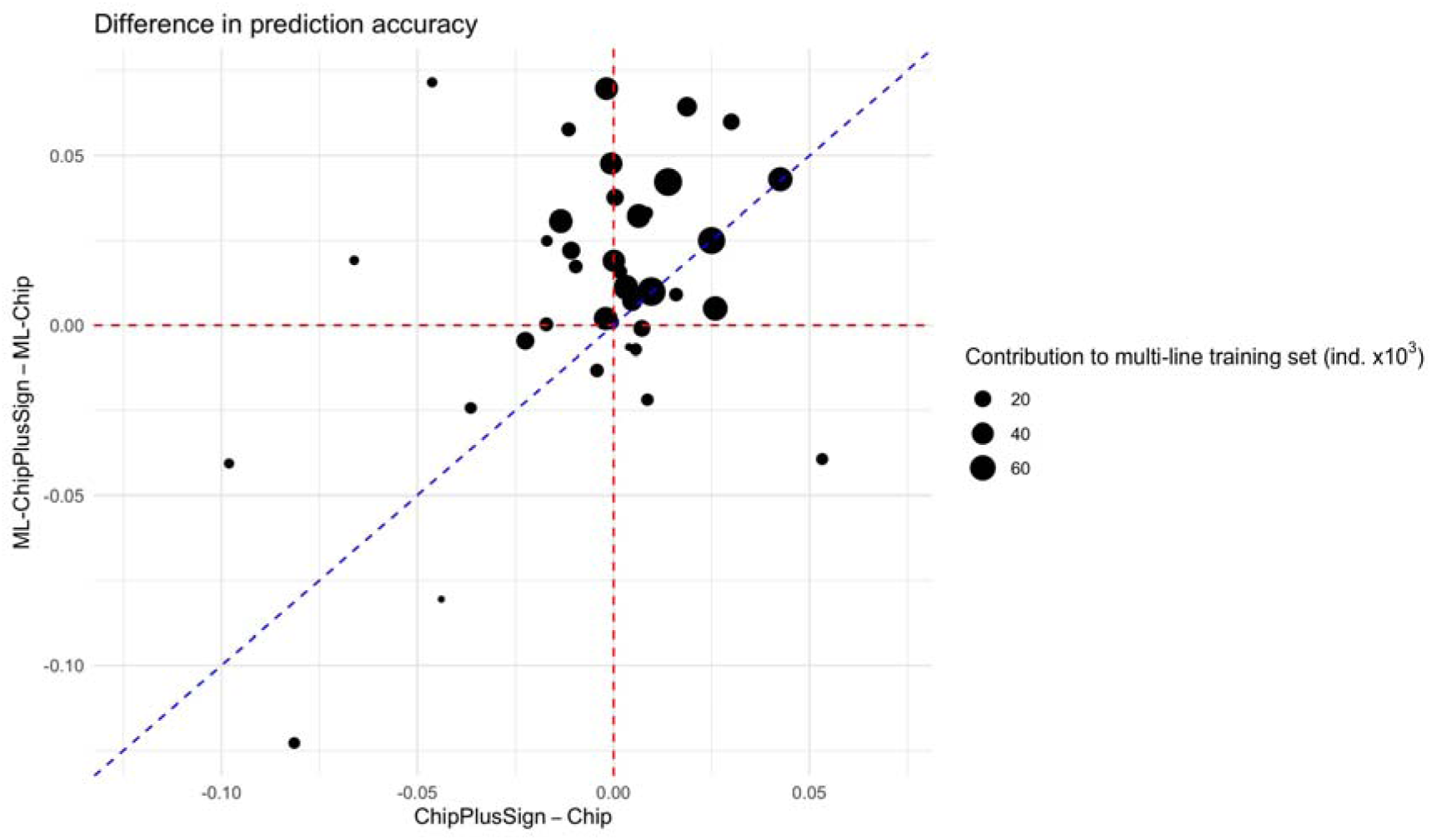
Comparison of the difference in genomic prediction accuracy in the multi-line scenarios (between ML-ChipPlusSign and ML-Chip) and in the within-line scenarios (between ChipPlusSign and Chip) for all traits and lines. Red dashed line at ‘no difference’. Blue dashed line is the bisector.

### Preselection of variants through genome-wide association study

Although GWAS with WGS has the potential to detect associations that are not captured by marker arrays, the fine-mapping of the associated regions and the preselection of variants through GWAS with WGS was limited due to the pervasiveness of linkage disequilibrium (Additional File 8) and was affected by false positives in a more severe way than GWAS with marker arrays, especially for highly polygenic traits (Additional File 4).

## Discussion

Our results showed that WGS has some potential to improve genomic prediction accuracy compared to marker arrays in intensely selected pig lines, but the use of WGS in current implementations should be carefully evaluated. On one hand, the small and non-robust improvements indicated that the strategies that we tested were likely suboptimal. On the other hand, the positive trend for the largest training sets indicated that we might have not reached the critical mass of data that is needed to leverage the potential of WGS, especially in scenarios where genomic prediction with marker arrays is already yielding high accuracy. The results from several traits and lines with different training set sizes allowed us to identify the most favourable scenarios for genomic prediction with WGS. We will discuss (1) the prediction accuracy that we achieved with WGS compared to commercial marker array data and the scenarios in which WGS may become beneficial, (2) the potential pitfalls for its effective implementation and the need for an optimised strategy, and (3) the suitability of WGS for genomic prediction.

### Prediction accuracy with whole-genome sequence data

We compared the genomic prediction accuracy of the current marker array (Chip) with sets of preselected WGS variants in a way that the number of variants remained similar across sets. Improvements in prediction accuracy can be limited if current marker arrays are already sufficiently dense to capture a large proportion of the genetic diversity in intensely selected livestock populations. These populations typically have small effective population size due to intense selective breeding [12,17]. Nevertheless, modest improvements have been achieved under certain scenarios across several studies. In our study, the most robust results were obtained with the ChipPlusSign variant sets, where the marker array was augmented with WGS variants that had statistically significant associations to the trait. This is consistent with previous reports that showed an improvement in prediction accuracy under similar approaches [28–31]. We augmented the marker array with 23 to 1083 significant variants in different scenarios. In the most successful scenarios, a minimum of around 200 significant variants were added and prediction accuracy improved by 0.025 on average with training sets of around 80k individuals. Other studies suggested additions of a larger number of variants. In Nordic cattle, adding 1623 variants (preselected as the combination of 3-5 variants for each of the top QTL per trait and breed) to a 50k marker array increased reliability (accuracy squared) by up to 0.05 [28], but a similar approach produced negligible improvements for low heritability traits [58]. In Holstein cattle, adding around 16k variants (preselected as the largest allele substitution effects) to a 60k marker array increased reliability on average by 0.027 (up to 0.048) [29]. In Hanwoo cattle, adding around 12k variants (3k for each of four traits) to a custom 50k marker array improved accuracy by up to ∼0.06 [31]. In sheep, adding around 400 variants (preselected by GWAS with regional heritability mapping) to a 50k marker array increased accuracy by 0.09 [30].

The modest performance of ChipPlusSign and Top40k could also be a consequence of the difficulty for fine-mapping causal variants through GWAS on WGS. Theoretically, the identification of all causal variants associated with a trait should improve genomic prediction accuracy [59]. Even though WGS allows the detection of a very large number of associations, problems such as false positives or p-value inflation also become more severe, so that the added noise might offset the detected signal. For instance, results in cattle showed that GWAS on WGS did not detect clearer associated regions relative to marker arrays and failed to capture QTL for genomic prediction [13], because the effect of potential QTL were spread across multiple variants. Therefore, WGS performed better with simple genetic architectures (i.e., traits with a low number of QTN). This is consistent with expectations and simulation results [60] that indicated that the benefit of WGS for genomic prediction would be limited by the number and size of QTN. Therefore, for largely polygenic traits (as most traits of interest in livestock production), training sets need to be very large before WGS can increase genomic prediction accuracy [60].

The advantage of using WGS might be limited by the small effective population size of livestock populations under selection [61] and by the current training set sizes, especially in scenarios where marker arrays are already yielding high genomic prediction accuracy [13,18]. Multi-line training sets could be particularly beneficial with the use of WGS because they allow a larger training set with low pairwise relationship degree among individuals. Previous simulations suggested that WGS might be the most beneficial with multi-breed reference panels [62], especially for numerically small populations. Our results with a multi-line training set indicated that WGS can improve prediction accuracy in scenarios that are less optimised than within-line genomic prediction by up to 0.04. However, in general those predictions were still less accurate than in within-line scenarios. In our multi-line scenarios, we only used variation that segregated across all seven lines. We observed that population-specific variation accounted only for small fractions of genetic variance [50] and it seems unlikely that they would contribute much to genomic prediction accuracy across breeds. Another possible obstacle is the differences in the allele substitution effects of the causal mutations across breeds. This can be caused by differences in allele frequency, contributions of non-additive effects and different genetic backgrounds, or even gene-by-environment interactions among others [22,63].

We observed low robustness of genomic prediction with WGS across traits and lines, and drops in prediction accuracy in some scenarios. Regarding bias, it has been noted that using the same reference individuals for preselecting variants through GWAS and for training the predictive equation can reduce genomic prediction accuracy and bias the predicted genomic breeding values [15,64]. In complementary tests, we observed no systematic increase in accuracy or bias after splitting the training set into two exclusive subsets, one for GWAS to preselect the predictor variants and the other for training the predictive equation (Additional File 9). One hypothesis is that both subsets belonged to the same population and therefore retained similar inter-relationships (i.e., they are not strictly independent sets of individuals). Moreover, the reduction in individuals available for training the predictors negatively affected genomic prediction accuracy.

We did not directly test persistence of genomic prediction accuracy across generations, but previous studies with empirical data found no higher persistence of prediction accuracy with WGS, not even with low degree of relationship between training and testing sets [13]. We expect such obstacles to persistence of accuracy until causal variants can be successfully identified and their non-additive effects are understood.

### Suboptimal strategy and pitfalls

The use of WGS for genomic prediction can only be reached after many other steps are completed to produce genotype data at the whole-genome level. Each of these steps has potential pitfalls to which the success of using WGS is sensitive. This strategy includes the choice of which individuals to sequence, the bioinformatics pipeline to call variants, the imputation of the WGS, and filtering of variants. When combined with the multiplicity of methods for preselecting variants for genomic prediction (which is unavoidable with current datasets, genomic prediction methods, and computational resources), there are many variables in the whole process that can affect the final result and that are not yet well understood. Therefore, a much greater effort for optimising such pipelines is required. Here we tested relatively simple approaches to evaluate how they performed with large WGS datasets. We have discussed what in our opinion are the main pitfalls of our approach for selection of the individuals to sequence [52] and the biases that may appear during processing of sequencing reads [47] elsewhere. Here we will focus discussion on imputation of WGS and its use for genomic prediction.

#### Imputation accuracy

Imputation of WGS is particularly challenging because typically we must impute a very large number of variants for a very large number of individuals from few sequenced individuals. As a consequence, genotype uncertainty can be high [19,25,65,66]. The accuracy of the imputed WGS is one of the main factors that may limit its potential for genomic prediction. In a simulation study, van den Berg et al. [25] quantified the impact of imputation errors on genomic prediction accuracy and showed that prediction accuracy decreases as errors accumulate, especially in the testing set.

We assessed the imputation accuracy of our approach elsewhere [38,52] and recommended that ∼2% of the population should be sequenced in intensely selected populations. In our study, line D was the line where genomic prediction accuracy with Top40k performed the worst, mostly performing worse than with the marker array. In this line, only 0.9% of the individuals in the population had been sequenced and therefore lower imputation accuracy could be expected. Although there was not enough evidence for establishing a link between these two features (sequencing effort and genomic prediction accuracy), we recommend cautious design of a sequencing strategy that is suited to the intended imputation method [52].

Genomic prediction accuracy could be improved by accounting for genotype uncertainty of the imputed WGS. For that, it could be advantageous to use allele dosages rather than best-guess genotypes [66], although most current implementations of genomic prediction methods cannot handle such information.

#### Preselection of predictor variants

Using WGS to simply increase the number of variants does not improve genomic prediction accuracy [16,19,22]. Due to the large number of variants in WGS, there is a need to remove uninformative variants [22,30,62,65,67]. We can expect variants that are causal or at least informative about the causal variants, which depends on their distance to the causal variants, to be the most predictive [68]. For this reason, variants that are in weak linkage disequilibrium with causal mutations have a ‘dilution’ effect, i.e., they add noise and limit prediction accuracy [22,30,67]. However, if too stringent filters are applied during preselection of predictor variants, there is a risk of removing true causal variants, and that would debilitate persistence of accuracy across generations and across populations [62,69]. For instance, the impact of removing variants with low minor allele frequency can vary depending on the minor allele frequency of the causal variants as well as the distance between preselected and causal variants [68]. Losing causal or informative variants would negatively affect multi-line or multi-breed prediction.

A popular strategy to preselect variants for the prediction model is based on association tests. Genome-wide association studies on WGS are expected to confirm associations that were already detected with marker arrays and identify novel associations (e.g., [35,70]). However, preliminary inspection of our empirical GWAS results showed that the added noise could easily offset the added information and fine-mapping remains challenging. Multi-breed GWAS [4] and meta-analyses [71] are suitable alternatives for GWAS to accommodate much larger population sizes and for combining results of populations with diverse genetic backgrounds. Multi-breed GWAS can be more efficient to identify informative variants than single-breed GWAS, which may benefit even prediction within lines [72]. Because the signal of some variants may go undetected for some traits but not for other correlated traits, combining GWAS information of several traits can also help identifying weak or moderate associations [23]. We did not test whether combining the significant markers from the different single-trait GWAS yielded greater improvements in prediction accuracy [28,31]. Multi-trait GWAS could be more suited for that purpose [70,73]. To improve fine-mapping, other GWAS models that incorporate biological information have been proposed (e.g., functional annotation [74] or metabolomics [75]).

Other methods were suggested to improve variant preselection for genomic prediction. VanRaden et al. [29] suggested that preselecting variants based on the genetic variance that they contribute rather than the significance of the association could be more advantageous, because the former would indirectly preselect variants with higher minor allele frequency. Other authors proposed preselection of variants using others statistics, such as the fixation index (F_ST_) between groups of individuals with high and low phenotype values to avoid the negative impact of spurious associations [67].

#### New models and methods

It is also likely that genomic prediction models, estimation methods, and their implementations need to be improved to leverage the potential of WGS. This is an active area of research and multiple novel methodologies have been proposed over the last years. Some examples are a combination of subsampling and Gibbs sampling [76], and a model that simultaneously fits a GBLUP term for a polygenic effect and a BayesC term for variants with large effects selected by the model (BayesGC) [24]. Testing alternative models and methods for genomic prediction was out of the scope of this study. However, together with refinements in the preselection of variants, it remains an interesting avenue for further optimisation of the analysis pipeline.

Some of the most promising methods are designed to incorporate prior biological information into the models. One of such methods is BayesRC [21], which extends BayesR by assigning flatter prior distributions to classes of variants that are more likely to be causal [17,20]. Similarly, GFBLUP [77] could be used to incorporate prior biological information from either QTL databases or GWAS as genomic features [19,34,65]. The model MBMG [26], which fits two genomic relationship matrices according to prior biological information, has also been proposed for multi-breed scenarios to improve genomic prediction in small populations. Haplotype-based models have been shown to provide greater prediction accuracy with WGS than variant-based models in pigs [78] and cattle [79]. However, the uptake of such models has been limited so far due to additional complexity, for example, to define haplotype blocks.

### Suitability of whole-genome sequence data for genomic prediction

The small improvements in genomic prediction accuracy that we achieved with WGS reflect the limited dimensionality of genomic information [61]. The WGS variants only produce small increases in prediction accuracy compared to marker arrays because the effective population size of intensely selected livestock populations is typically small and marker arrays already capture a large proportion of their independent chromosome segments. Thus, the use of WGS in current implementations of genomic prediction should be carefully evaluated against the cost of generating the WGS, especially given the large size of the datasets that are required. Sequencing costs are expected to continue to decrease and therefore large datasets of WGS will become more affordable in time, while efforts to develop and optimise scalable and accurate pipelines for WGS-based data generation, storage, and analysis are on-going (e.g., [80,81]). These advances, together with a finer knowledge of the genetic architecture of traits empowered by WGS, could allow a case-by-case refinement of genomic prediction. However, to date, the low robustness of the results for complex traits discourage the generalised use of WGS for traits that are already accurately predicted by conventional means.

## Conclusion

Our results showed that WGS has some potential to improve genomic prediction accuracy compared to marker arrays in intensely selected pig lines. However, the prediction accuracy with each set of preselected WGS variants was not robust across traits and lines and the improvements in prediction accuracy that we achieved so far with WGS compared to marker arrays were generally small. The most favourable results for WGS were obtained when the largest training sets were available and used to preselect variants with statistically significant associations to the trait for augmenting the established marker array. With this method and training sets of around 80k individuals, average improvements of genomic prediction accuracy of 0.025 were observed in within-line scenarios. A combination of larger training sets and improved pipelines could further improve genomic prediction accuracy. The robustness of the whole strategy for generating WGS at the population level must be carefully stress-tested and further optimised. However, with the current implementations of genomic prediction, the use of WGS should be carefully evaluated on a case-by-case basis against the cost of generating the WGS at a large scale.

## Ethics approval and consent to participate

The samples used in this study were derived from the routine breeding activities of PIC.

## Consent for publication

Not applicable.

## Availability of data and material

The software packages AlphaPhase, AlphaImpute, and AlphaPeel are available from https://github.com/AlphaGenes. The software package AlphaSeqOpt is available from the AlphaGenes website (http://www.alphagenes.roslin.ed.ac.uk). The datasets generated and analysed in this study are derived from the PIC breeding programme and not publicly available.

## Authors’ contributions

RRF, GG, and JMH designed the study; CYC assisted in preparing the datasets; RRF, AW and MJ performed the analyses; RRF wrote the first draft; AW, CYC, BDV, WHO, GG, and JMH assisted in the interpretation of the results and provided comments on the manuscript. All authors read and approved the final manuscript.

## Acknowledgements

This work has made use of the resources provided by the Edinburgh Compute and Data Facility (ECDF) (http://www.ecdf.ed.ac.uk/).

