## Additional File 1 for "Genomic prediction with whole-genome sequence data in intensely selected pig lines"

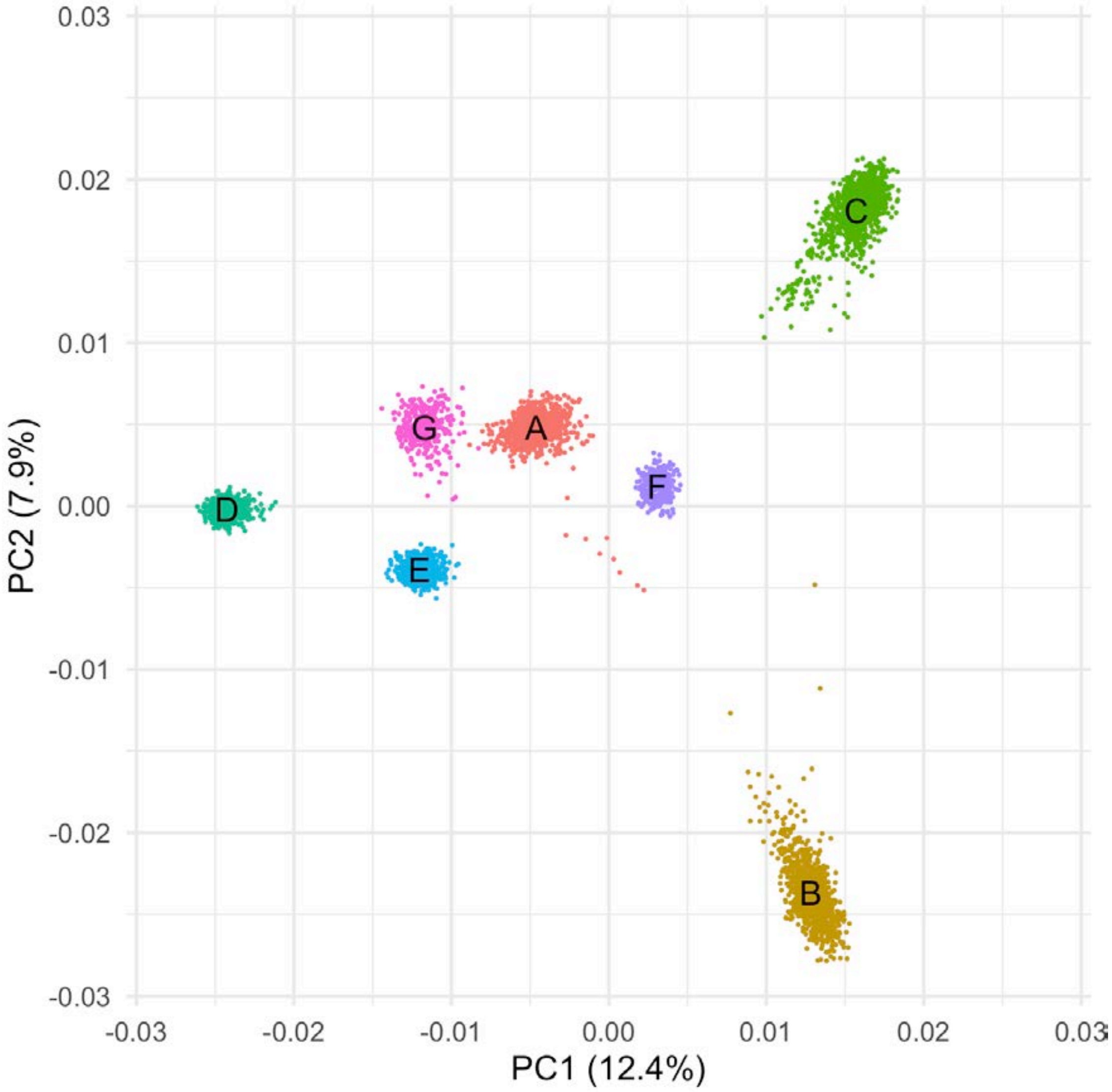


**Figure S1.** Population structure of the sequenced pigs according to the two first principal components. The colour clusters correspond to lines A to G.
