## Additional File 2 for "Genomic prediction with whole-genome sequence data in intensely selected pig lines"

### Prediction accuracy for all traits and lines

Left: Correlation. Dashed line at value of Chip as a reference. Values indicate relative difference to reference Chip. Right: Bias. Dashed line at the ideal value.

<u>ADG</u>

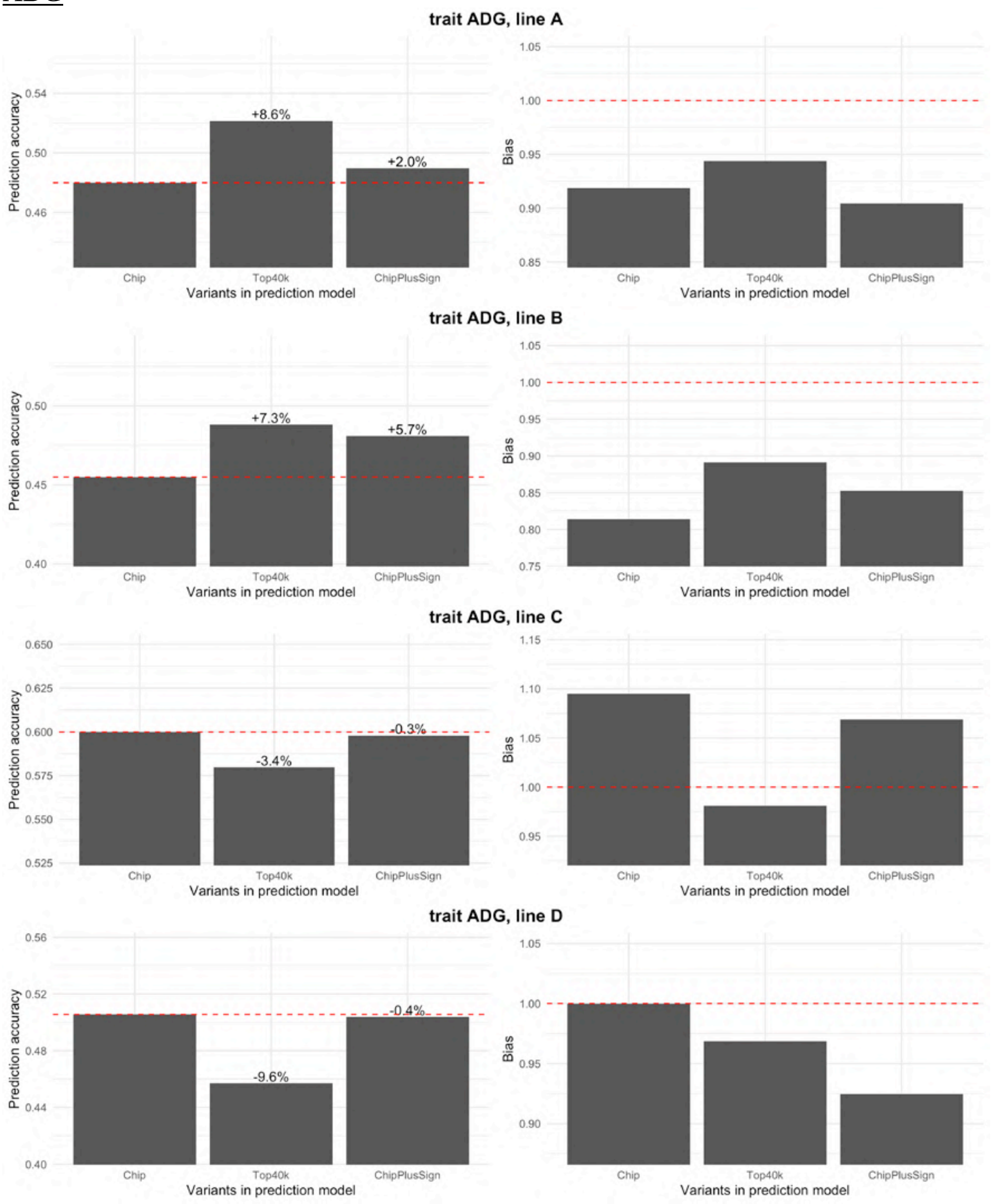

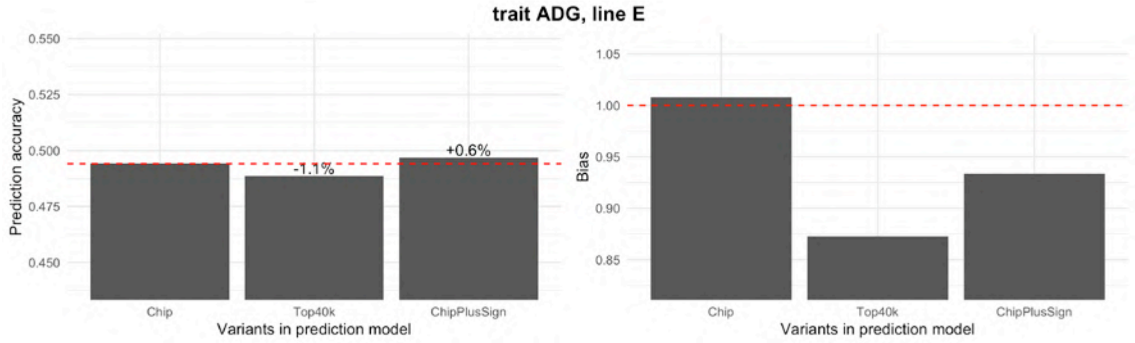
trait ADG, line E
Prediction accuracy
Bias
+0.6%
-1.1%
Chip
Top40k
ChipPlusSign
Variants in prediction model

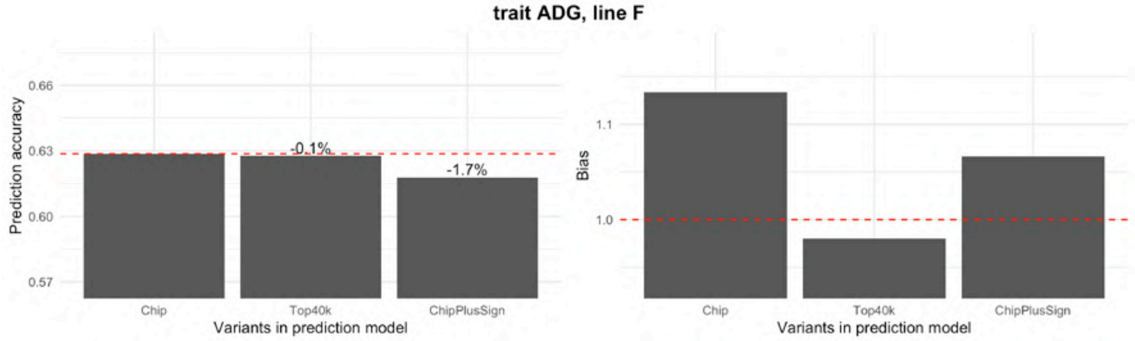
trait ADG, line F
Prediction accuracy
Bias
-0.1%
-1.7%
Chip
Top40k
ChipPlusSign
Variants in prediction model

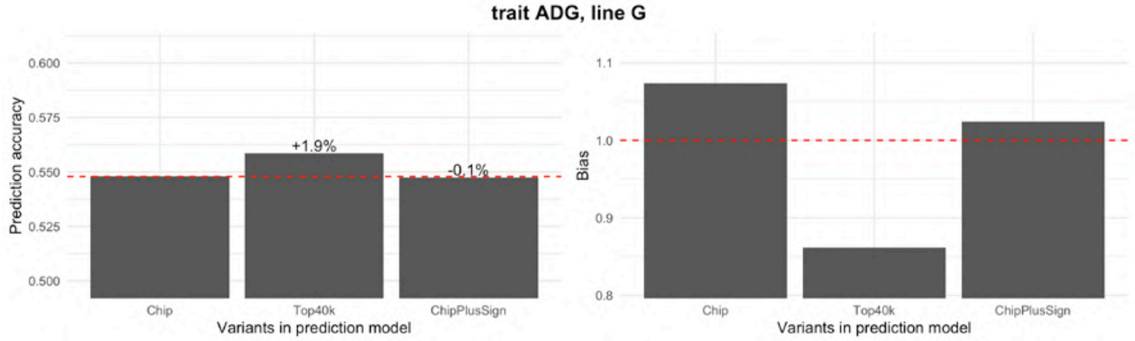
trait ADG, line G
Prediction accuracy
Bias
+1.9%
-0.1%
Chip
Top40k
ChipPlusSign
Variants in prediction model

<u>BFT</u>

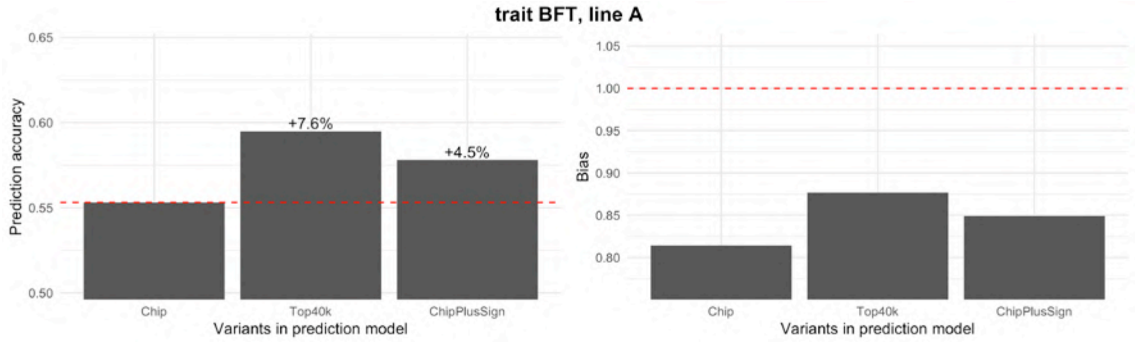
trait BFT, line A
Prediction accuracy
Bias
+7.6%
+4.5%
Chip
Top40k
ChipPlusSign
Variants in prediction model

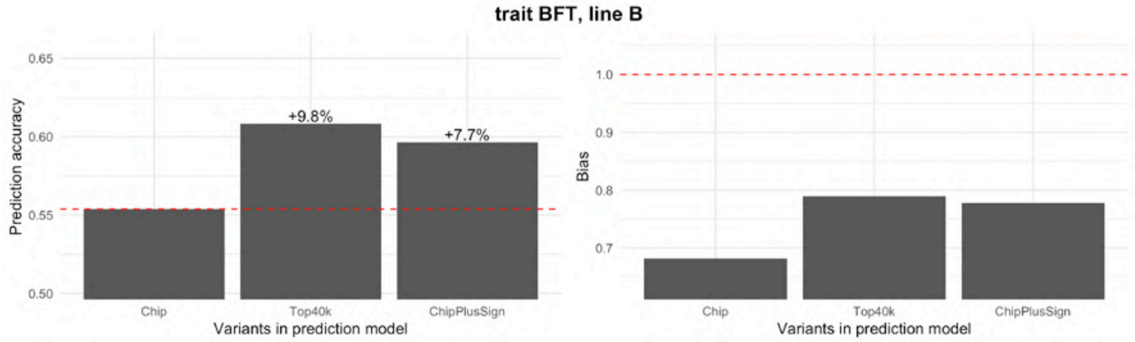
trait BFT, line B
Prediction accuracy
Bias
+9.8%
+7.7%
Chip
Top40k
ChipPlusSign
Variants in prediction model

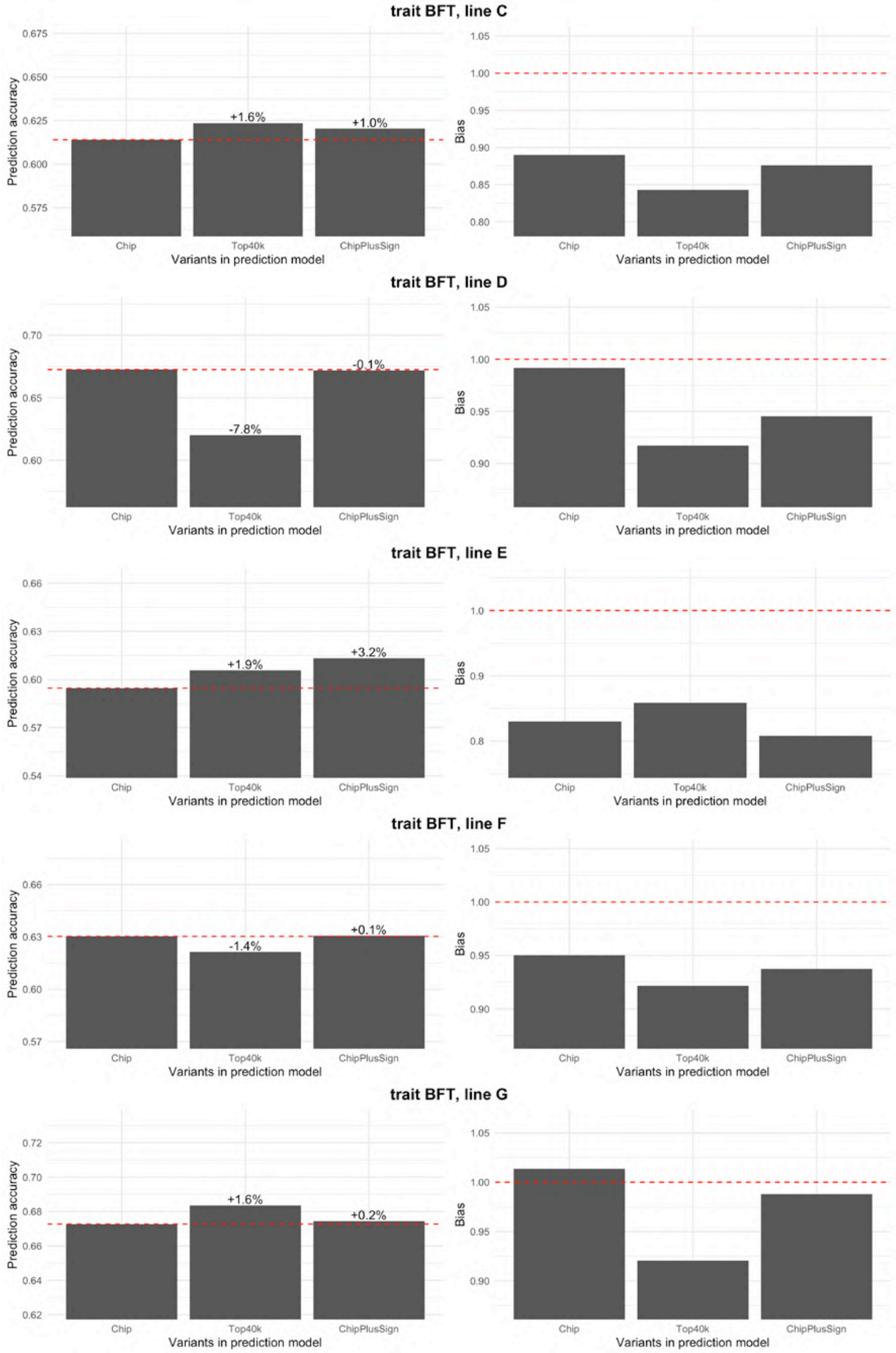
trait BFT, line C
Prediction accuracy
0.675
0.650
0.625
0.600
0.575
+1.6%
+1.0%
Chip
Top40k
ChipPlusSign
Variants in prediction model
Bias
1.05
1.00
0.95
0.90
0.85
0.80
Chip
Top40k
ChipPlusSign
Variants in prediction model
trait BFT, line D
Prediction accuracy
0.70
0.65
0.60
-0.1%
-7.8%
Chip
Top40k
ChipPlusSign
Variants in prediction model
Bias
1.05
1.00
0.95
0.90
Chip
Top40k
ChipPlusSign
Variants in prediction model
trait BFT, line E
Prediction accuracy
0.66
0.63
0.60
0.57
0.54
+1.9%
+3.2%
Chip
Top40k
ChipPlusSign
Variants in prediction model
Bias
1.0
0.9
0.8
Chip
Top40k
ChipPlusSign
Variants in prediction model
trait BFT, line F
Prediction accuracy
0.66
0.63
0.60
0.57
-1.4%
+0.1%
Chip
Top40k
ChipPlusSign
Variants in prediction model
Bias
1.05
1.00
0.95
0.90
Chip
Top40k
ChipPlusSign
Variants in prediction model
trait BFT, line G
Prediction accuracy
0.72
0.70
0.68
0.66
0.64
0.62
+1.6%
+0.2%
Chip
Top40k
ChipPlusSign
Variants in prediction model
Bias
1.05
1.00
0.95
0.90
Chip
Top40k
ChipPlusSign
Variants in prediction model

LD

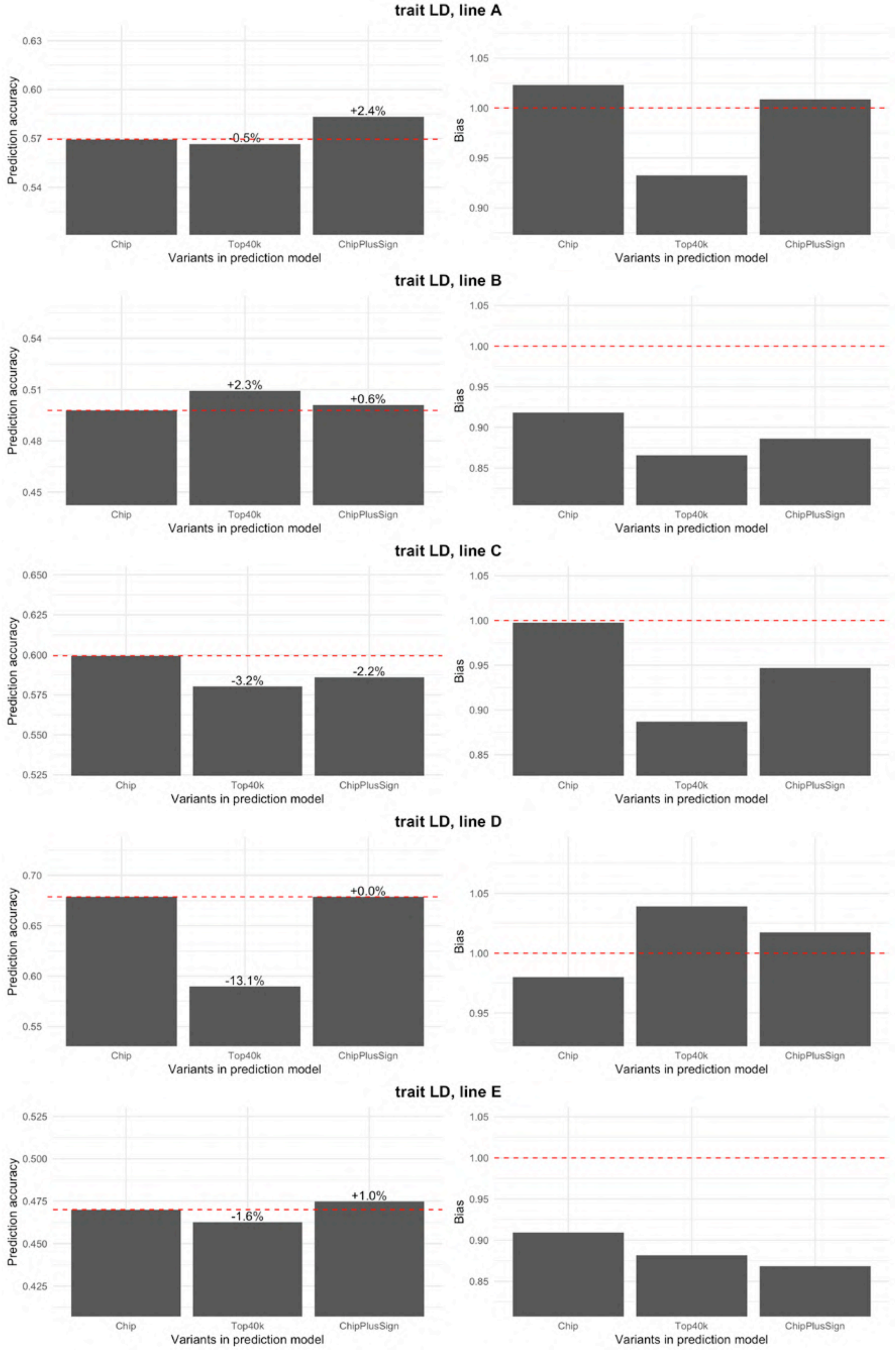
trait LD, line A
Prediction accuracy
0.63
0.60
0.57
0.54
+2.4%
-0.5%
Bias
1.05
1.00
0.95
0.90
Chip
Top40k
ChipPlusSign
Variants in prediction model
trait LD, line B
0.54
0.51
0.48
0.45
+2.3%
+0.6%
1.05
1.00
0.95
0.90
0.85
trait LD, line C
0.650
0.625
0.600
0.575
0.550
0.525
-3.2%
-2.2%
1.05
1.00
0.95
0.90
0.85
trait LD, line D
0.70
0.65
0.60
0.55
+0.0%
-13.1%
1.05
1.00
0.95
trait LD, line E
0.525
0.500
0.475
0.450
0.425
+1.0%
-1.6%
1.05
1.00
0.95
0.90
0.85

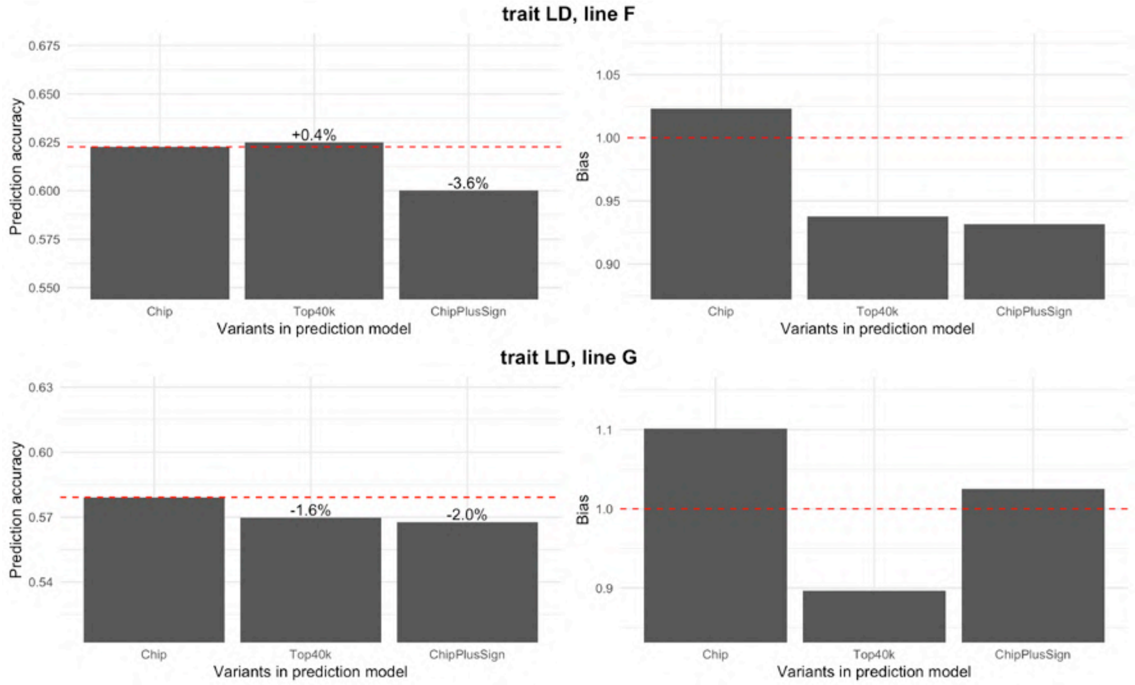
trait LD, line F
Prediction accuracy
+0.4%
-3.6%
Bias
Chip
Top40k
ChipPlusSign
Variants in prediction model
trait LD, line G
-1.6%
-2.0%

ADFI

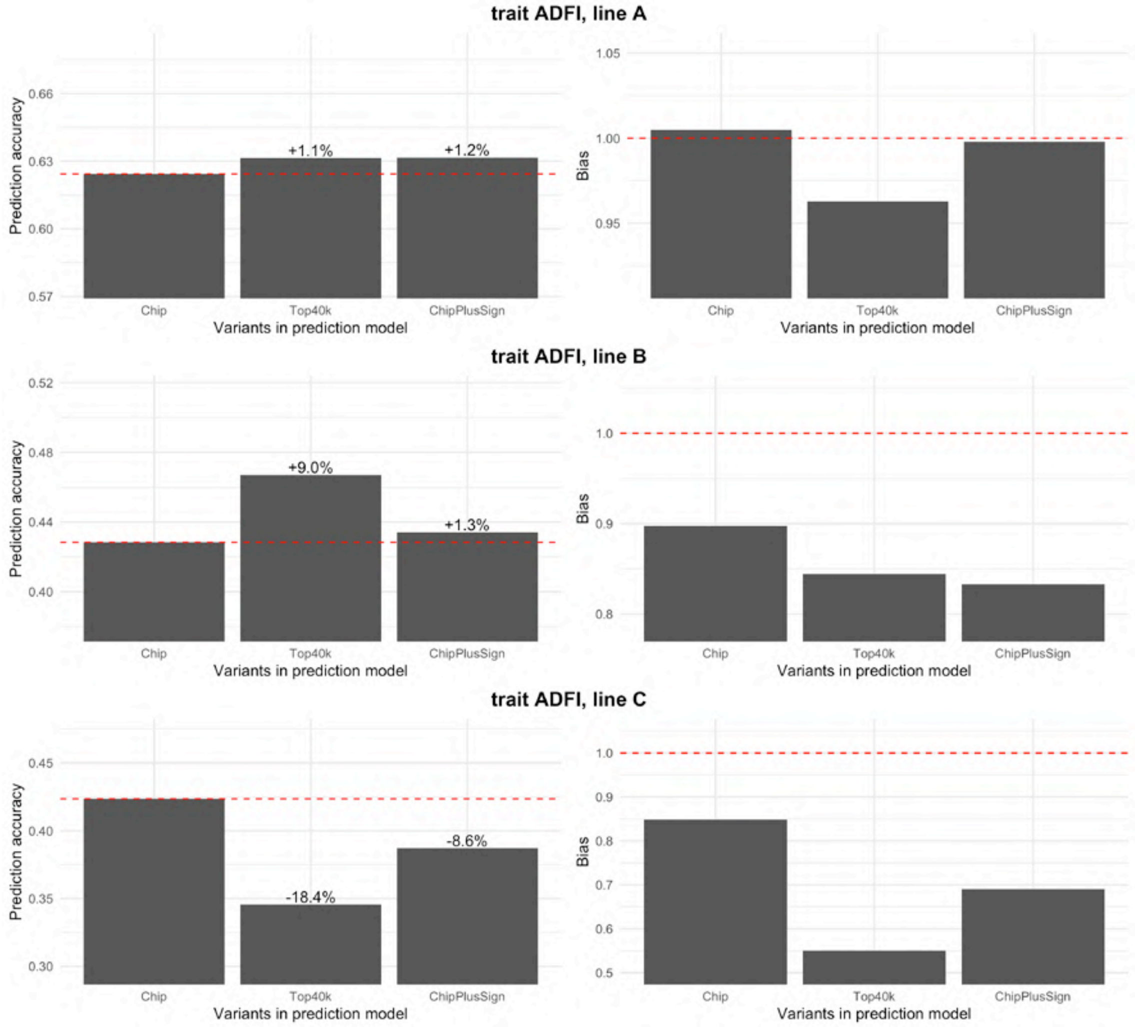
trait ADFI, line A
Prediction accuracy
+1.1%
+1.2%
Bias
Chip
Top40k
ChipPlusSign
Variants in prediction model
trait ADFI, line B
+9.0%
+1.3%
trait ADFI, line C
-18.4%
-8.6%

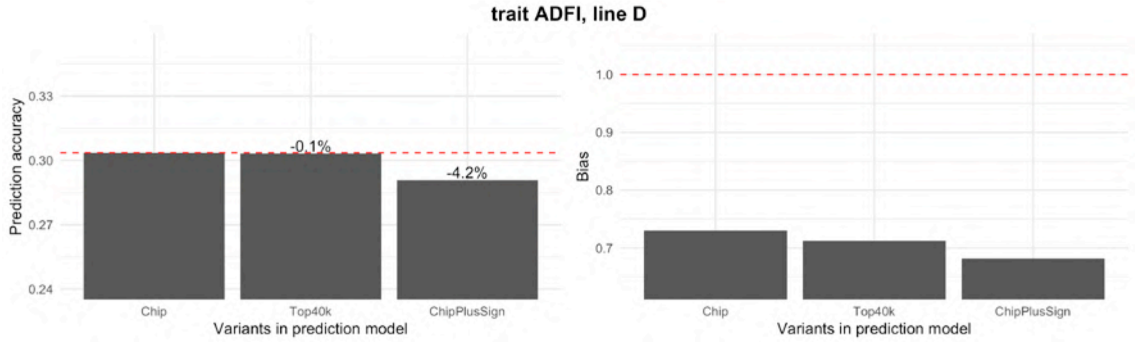
trait ADFI, line D
Prediction accuracy
0.33
0.30
0.27
0.24
-0.1%
-4.2%
Chip
Top40k
ChipPlusSign
Variants in prediction model
Bias
1.0
0.9
0.8
0.7
Chip
Top40k
ChipPlusSign
Variants in prediction model

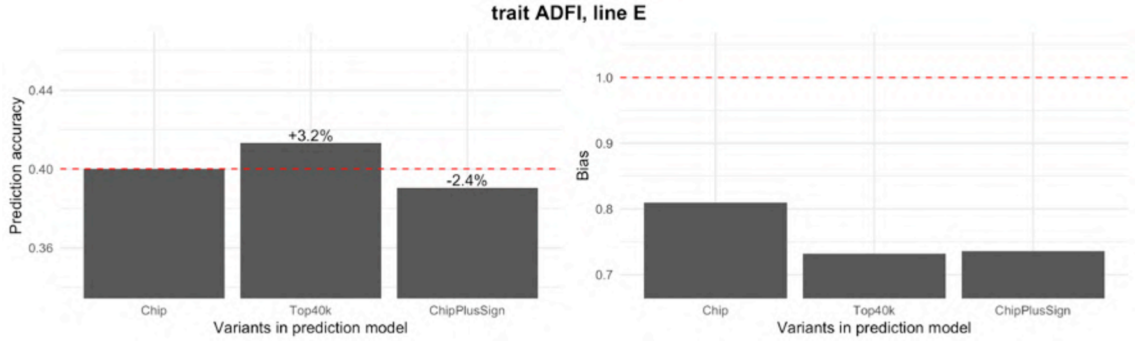
trait ADFI, line E
Prediction accuracy
0.44
0.40
0.36
+3.2%
-2.4%
Chip
Top40k
ChipPlusSign
Variants in prediction model
Bias
1.0
0.9
0.8
0.7
Chip
Top40k
ChipPlusSign
Variants in prediction model

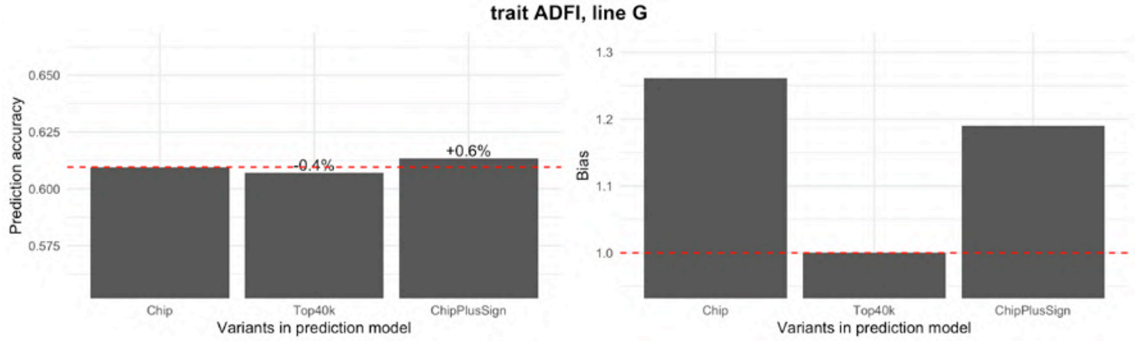
trait ADFI, line G
Prediction accuracy
0.650
0.625
0.600
0.575
-0.4%
+0.6%
Chip
Top40k
ChipPlusSign
Variants in prediction model
Bias
1.3
1.2
1.1
1.0
Chip
Top40k
ChipPlusSign
Variants in prediction model

<u>FCR</u>

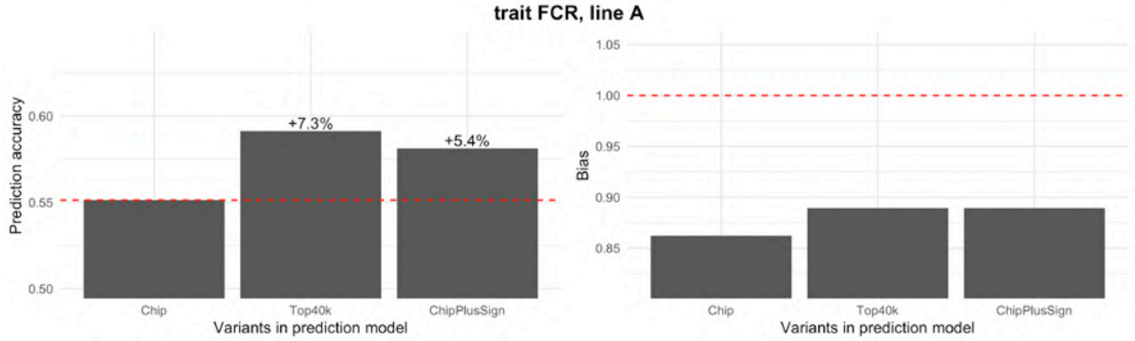
trait FCR, line A
Prediction accuracy
0.60
0.55
0.50
+7.3%
+5.4%
Chip
Top40k
ChipPlusSign
Variants in prediction model
Bias
1.05
1.00
0.95
0.90
0.85
Chip
Top40k
ChipPlusSign
Variants in prediction model

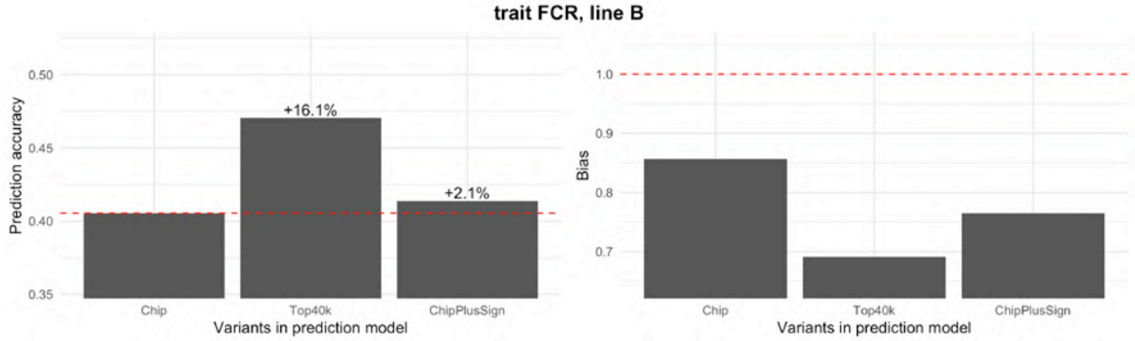
trait FCR, line B
Prediction accuracy
0.50
0.45
0.40
0.35
+16.1%
+2.1%
Chip
Top40k
ChipPlusSign
Variants in prediction model
Bias
1.0
0.9
0.8
0.7
Chip
Top40k
ChipPlusSign
Variants in prediction model

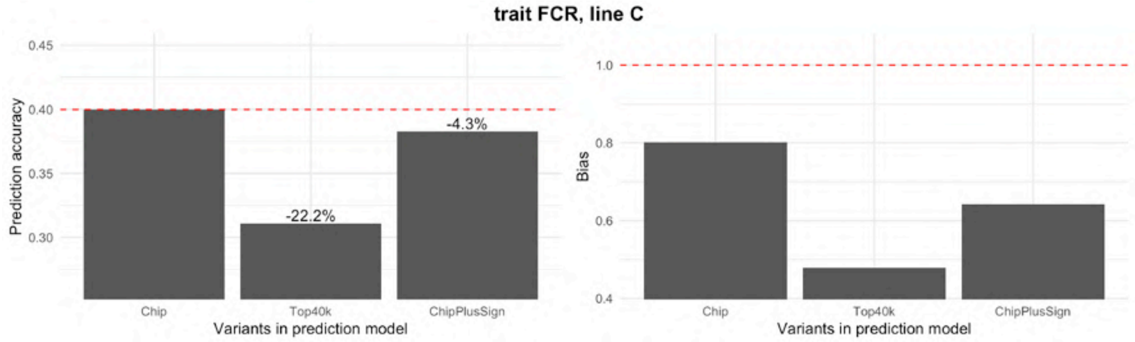
trait FCR, line C
Prediction accuracy
Bias
-22.2%
-4.3%
Chip
Top40k
ChipPlusSign
Variants in prediction model

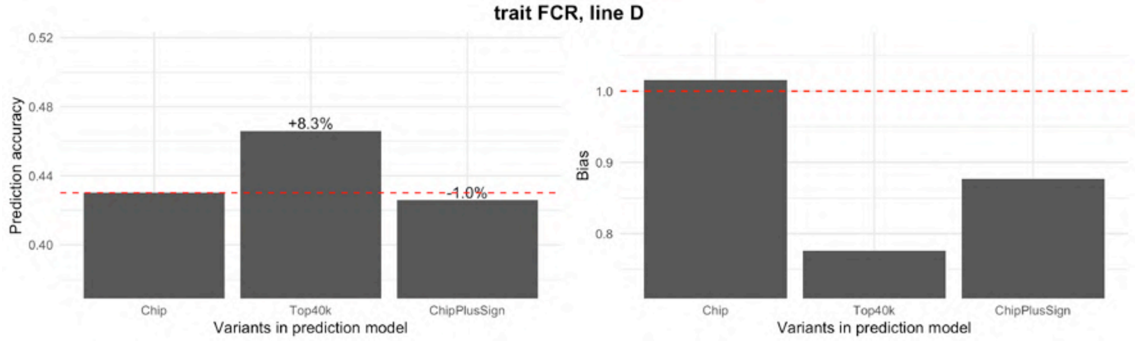
trait FCR, line D
Prediction accuracy
Bias
+8.3%
-1.0%
Chip
Top40k
ChipPlusSign
Variants in prediction model

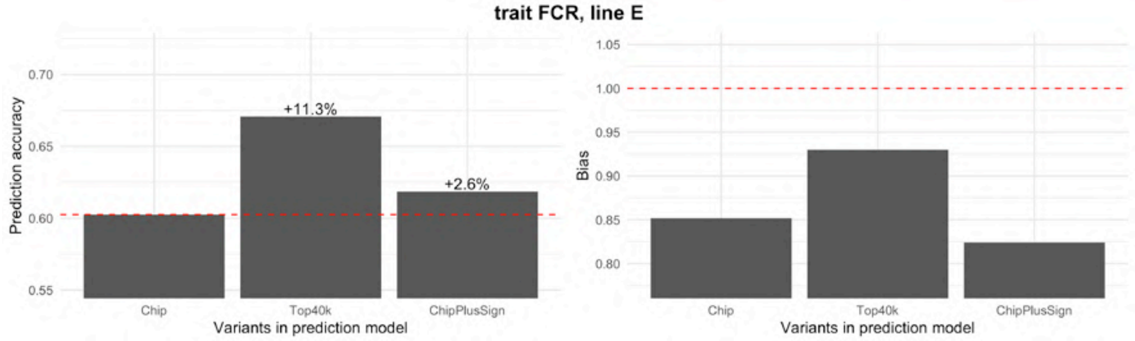
trait FCR, line E
Prediction accuracy
Bias
+11.3%
+2.6%
Chip
Top40k
ChipPlusSign
Variants in prediction model

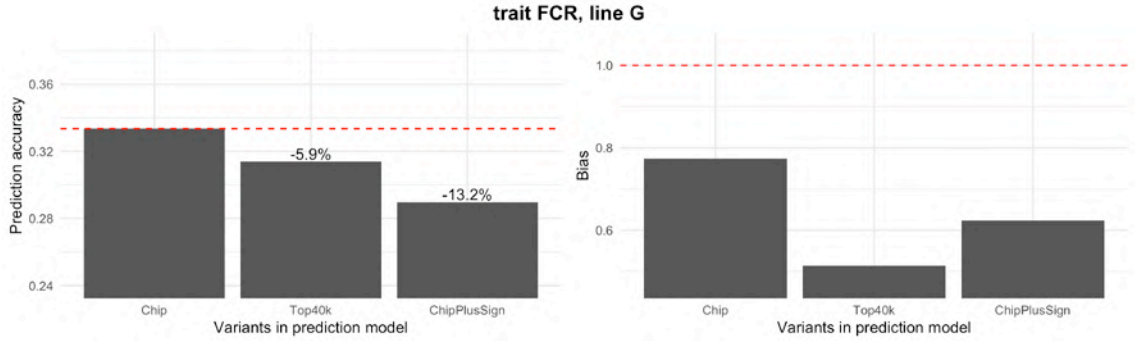
trait FCR, line G
Prediction accuracy
Bias
-5.9%
-13.2%
Chip
Top40k
ChipPlusSign
Variants in prediction model

TNB

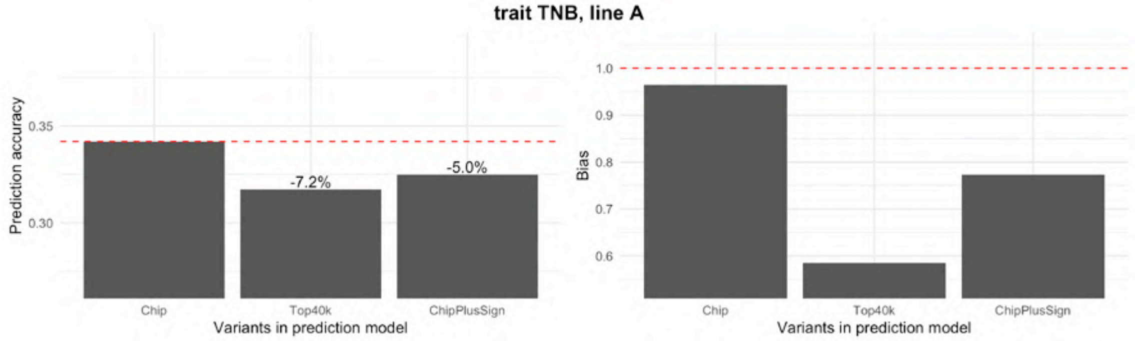
trait TNB, line A
Prediction accuracy
Bias
-7.2%
-5.0%
Chip
Top40k
ChipPlusSign
Variants in prediction model

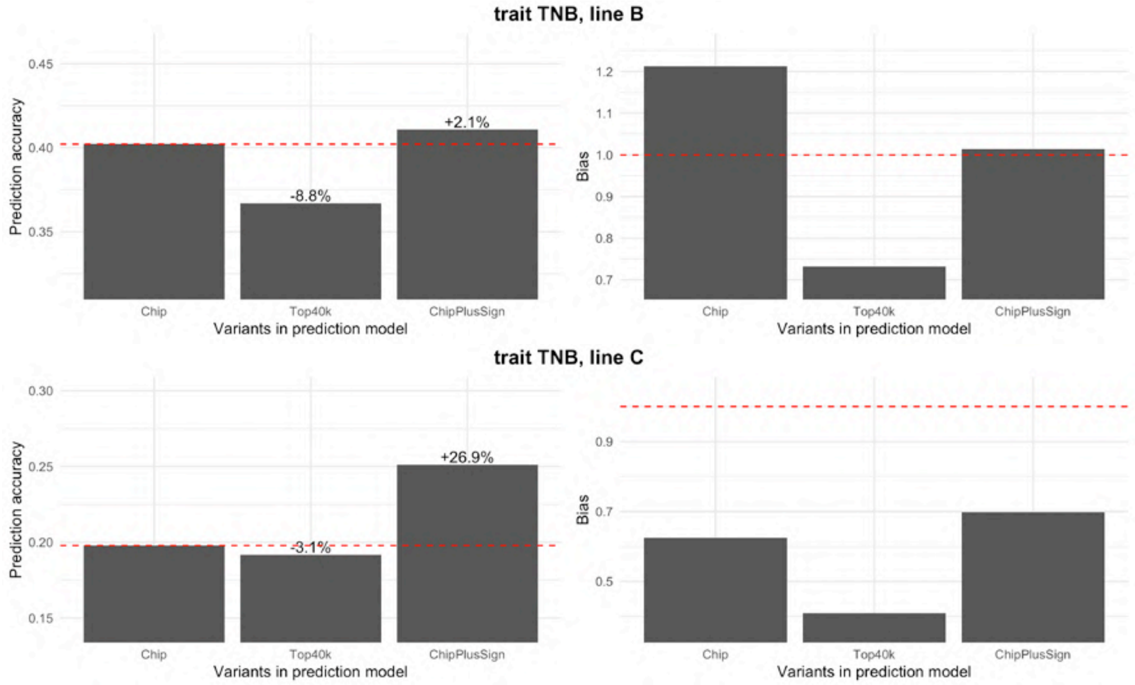
trait TNB, line B
Prediction accuracy
+2.1%
-8.8%
Bias
Chip
Top40k
ChipPlusSign
Variants in prediction model
trait TNB, line C
+26.9%
-3.1%

<u>LWW</u>

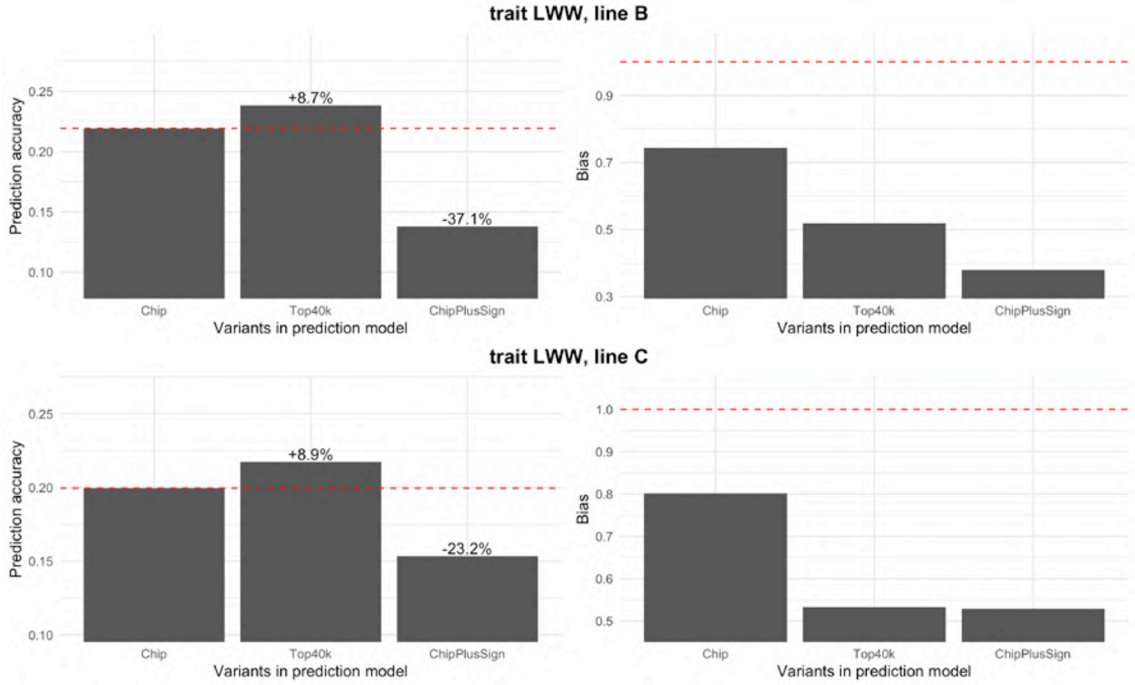
trait LWW, line B
Prediction accuracy
+8.7%
-37.1%
Bias
Chip
Top40k
ChipPlusSign
Variants in prediction model
trait LWW, line C
+8.9%
-23.2%

RET

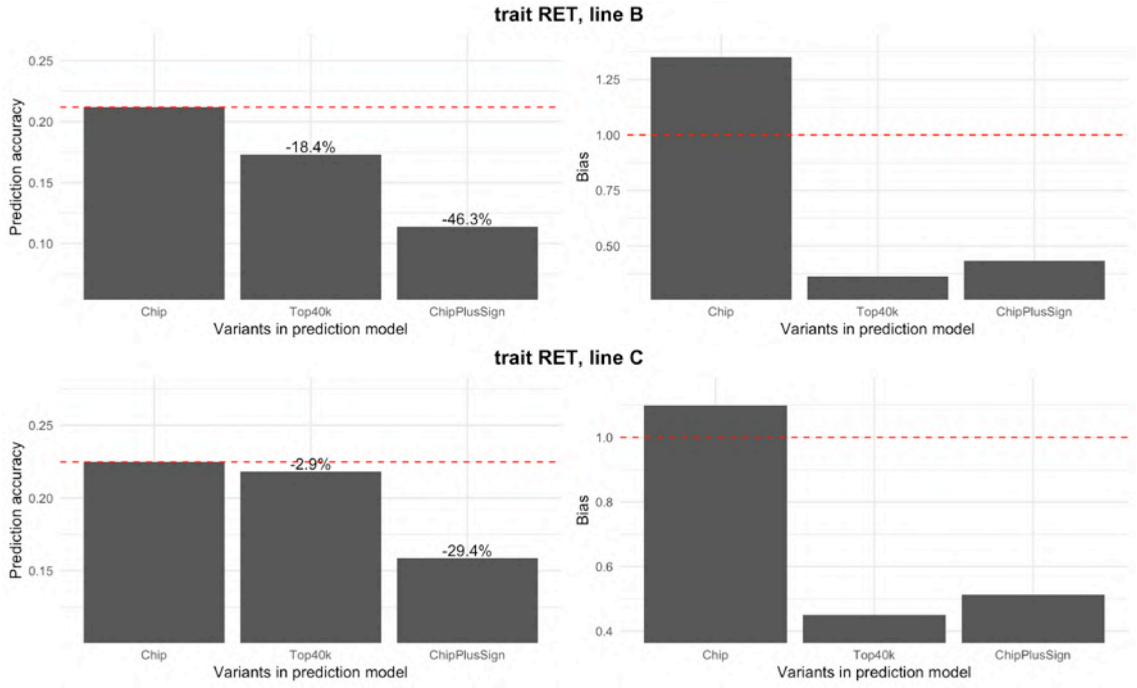
trait RET, line B
Prediction accuracy
-18.4%
-46.3%
Bias
Chip
Top40k
ChipPlusSign
Variants in prediction model
trait RET, line C
-2.9%
-29.4%
