## Additional File 3 for "Genomic prediction with whole-genome sequence data in intensely selected pig lines"

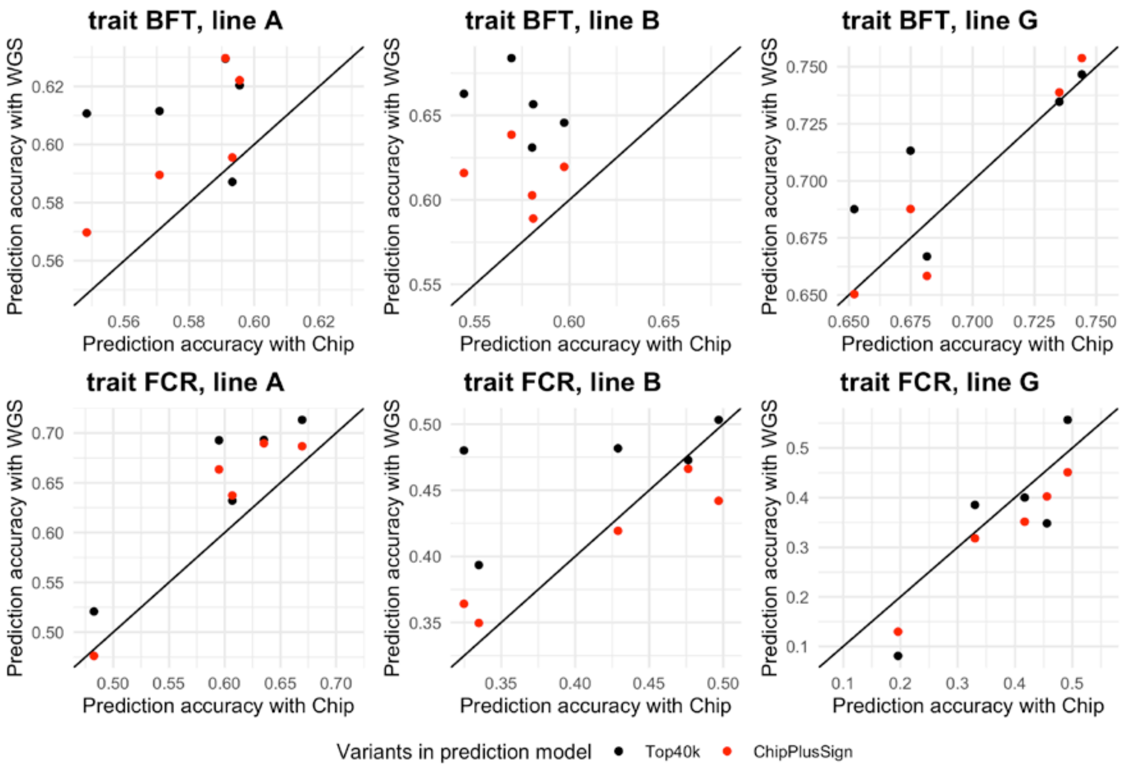


**Figure S2.** Genomic prediction accuracy obtained with each set of preselected whole-genome sequence (WGS) variants (either Top40k or ChipPlusSign) against that obtained with the marker array (Chip) in each of the five repetitions.

**Table S1.** Comparison of the results obtained with a single repetition or with five repetitions. The difference of the genomic prediction accuracy between each set of preselected whole-genome sequence variants (either Top40k or ChipPlusSign) and the marker array (Chip) is reported.

| Trait | Line | Top40k – Chip | | | ChipPlusSign – Chip | | |
|---|---|---|---|---|---|---|---|
| | | Single repetition | Five repetitions | | Single repetition | Five repetitions | |
| | | | Mean | SD | | Mean | SD |
| BFT | A | 0.042 | 0.032 | 0.025 | 0.025 | 0.021 | 0.013 |
| BFT | B | 0.054 | 0.082 | 0.034 | 0.043 | 0.039 | 0.030 |
| BFT | G | 0.011 | 0.012 | 0.023 | 0.002 | 0.000 | 0.014 |
| FCR | A | 0.040 | 0.053 | 0.028 | 0.030 | 0.033 | 0.030 |
| FCR | B | 0.065 | 0.054 | 0.063 | 0.009 | -0.004 | 0.035 |
| FCR | G | -0.020 | -0.024 | 0.086 | -0.044 | -0.048 | 0.022 |
