## Additional File 4 for "Genomic prediction with whole-genome sequence data in intensely selected pig lines"

### Genomic prediction and genome-wide association study using simulated traits

#### Methods

To assist the interpretation of the empirical results, we simulated nine traits with different numbers of quantitative trait nucleotides (QTN; 100, 1,000 or 10,000 QTN) and heritability levels ($h^2$; 0.10, 0.25 or 0.50). Positions of the QTN were sampled randomly amongst all variants called across all lines. Because QTN were sampled from all variants, some QTN were fixed in some of the lines while segregating in others. There were only negligible differences in the number of segregating QTN per line (53 to 61, 531 to 583, or 5375 to 6058, respectively). Effects of the QTN were sampled from a gamma distribution with shape of 2 and scale of 5. We calculated the breeding value for each individual using these QTN genotypes and effects. We then added a residual term sampled from a normal distribution with a variance parameter adjusted to produce a phenotype with the targeted heritability. A phenotype value was calculated for every individual with imputed genotypes that passed the imputation accuracy control (104,661 to 17,224 individuals per line; see Table 1 in the main text). In these simulations, we used the imputed genotypes as real genotypes and, therefore, implicitly cancelled any errors that might arise from the processing of the sequencing reads and genotype imputation.

Further analyses including GWAS and genomic prediction were performed as described in the main text for the empirical traits. Because of the high computational demands of these analyses, no repetitions were performed and therefore interpretation of these results should be cautious.

#### Results

##### Genomic prediction

The improvements of genomic prediction accuracy of whole-genome sequence data (WGS) compared to the marker array depended on the genetic architecture of the traits. Traits with high heritability and low number of QTN were more likely to show larger improvements in prediction accuracy. With Top40k (Figure S4.1), heritability seemed to be the main factor that affected the expected improvement with large training sets (from null improvements when $h^2$=0.1 to improvements of approximately 0.05 when $h^2$=0.5, regardless of the number of QTN, with a training set of 92k individuals). With ChipPlusSign (Figure S4.2), the expected improvements with the same training set (92k individuals) were not only greater in magnitude but depended on both heritability and the number of QTN (from null improvements when $h^2$=0.1 to improvements of approximately 0.03 to 0.10 when $h^2$=0.5 with a number of 100 to 10k QTN, respectively). Results confirmed the trends observed for the empirical traits; for instance, the higher robustness of ChipPlusSign compared to Top40k.

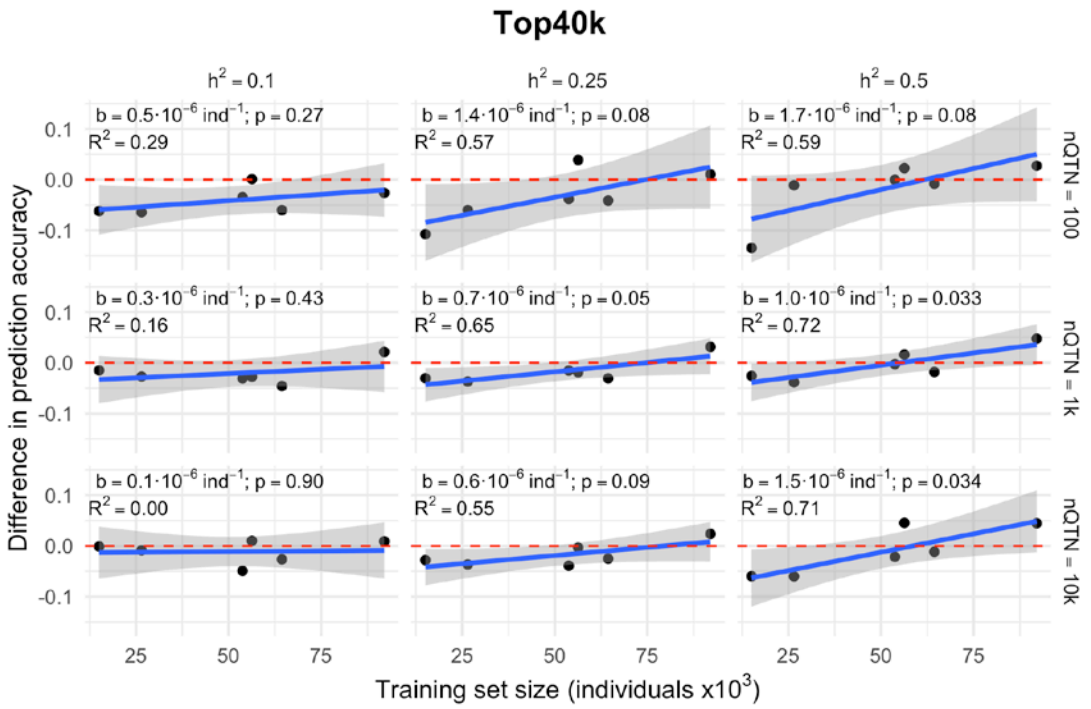


**Figure S4.1.** Genomic prediction accuracy with the Top40k variants for the simulated traits. The difference between the Top40k and marker array is shown by heritability ($h^2$) and number of quantitative trait nucleotides (nQTN) of the simulated traits. Red dashed line at 'no difference'. Regression coefficient (b) and p-value of training set size is provided, as well as the coefficient of determination ($R^2$) of the model.

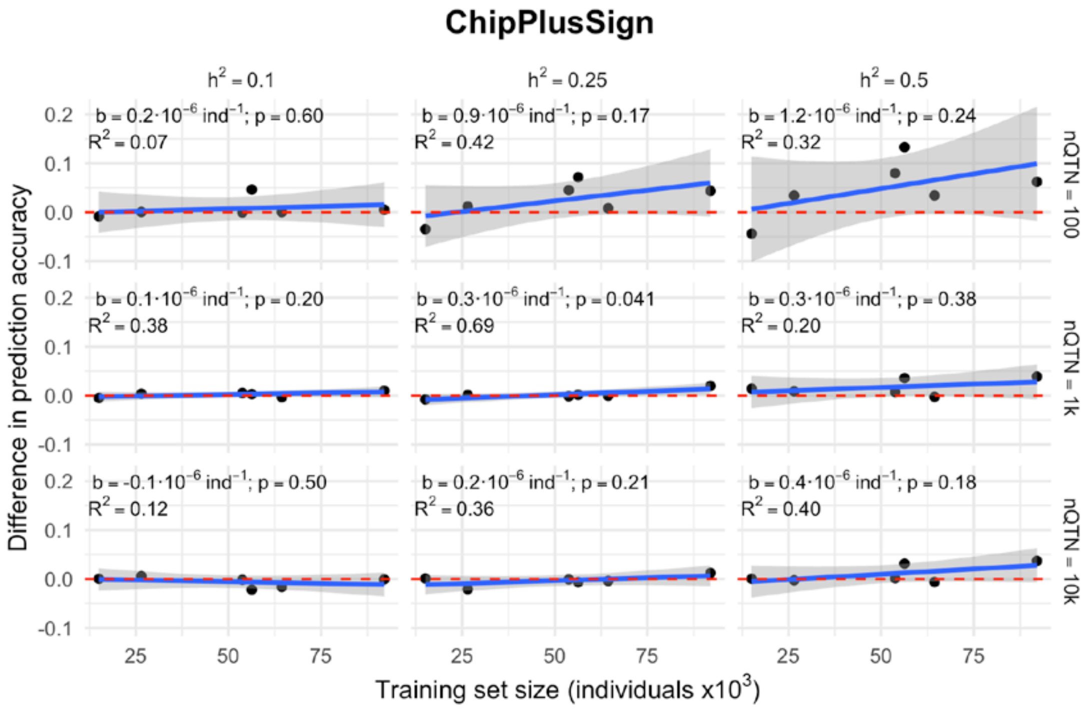


**Figure S4.2.** Genomic prediction accuracy with the ChipPlusSign variants for the simulated traits. The difference between the ChipPlusSign and marker array is shown by heritability ($h^2$) and number of quantitative trait nucleotides (nQTN) of the simulated traits. Other details as in Figure S4.1.

Genome-wide association study

We defined genomic regions that contained significant associations by overlapping 500-kb segments centred on the significant markers ($p \leq 10^{-6}$). Table S4.1 shows the number of regions with significant associations that were detected using either the marker array or WGS, and whether they contained zero, one or multiple QTN. The WGS detected a much larger proportion of QTN than the marker array, especially for the traits with high heritability and with large population sizes. The most favourable scenarios for identifying regions that contained unequivocally a single QTN with WGS were those in which the trait was controlled by a low number of QTN. However, even though the genetic architecture was very simple and consisted only of additive effects, the regions with significant associations only captured a small fraction of the QTN that segregated within each line. Moreover, using WGS also increased the number of regions with significant associations that contained no QTN, which could therefore be considered as false positives. Some of the selected regions contained multiple QTN, which could indicate either a 'hit by chance' or an inability to disentangle multiple causal variants. While false positives also occur with marker array, their incidence was more severe with the WGS, especially for traits with a large number of QTN. Large population sizes further aggravated the inflation of genome-wide p-values.

**Table S4.1.** Number of genomic regions around significantly associated markers that contained 0, 1 or 2 or more quantitative trait nucleotides (QTN).

| **$h^2$** | **nQTN** | **Line size** | **Marker array** | | **Whole-genome sequence** | | |
|---|---|---|---|---|---|---|---|
| | | | **0 QTN** | **1 QTN** | **0 QTN** | **1 QTN** | **≥2 QTN** |
| 0.10 | 100 | 27k | 4 | 1 | 8 | 6 | 0 |
| | | 56k | 11 | 3 | 19 | 19 | 0 |
| | | 92k | 10 | 7 | 44 | 19 | 0 |
| | 1k | 27k | 1 | 0 | 4 | 0 | 1 |
| | | 56k | 1 | 0 | 16 | 3 | 1 |
| | | 92k | 1 | 0 | 283 | 9 | 0 |
| | 10k | 27k | 1 | 0 | 1 | 0 | 0 |
| | | 56k | 0 | 0 | 16 | 2 | 1 |
| | | 92k | 2 | 0 | 186 | 17 | 12 |
| 0.25 | 100 | 27k | 11 | 6 | 26 | 15 | 1 |
| | | 56k | 22 | 8 | 44 | 28 | 3 |
| | | 92k | 20 | 7 | 90 | 34 | 1 |
| | 1k | 27k | 0 | 0 | 8 | 1 | 3 |
| | | 56k | 3 | 0 | 34 | 15 | 6 |
| | | 92k | 6 | 0 | 692 | 49 | 16 |
| | 10k | 27k | 0 | 0 | 2 | 0 | 0 |
| | | 56k | 0 | 0 | 90 | 9 | 22 |
| | | 92k | 4 | 0 | 564 | 56 | 164 |
| 0.50 | 100 | 27k | 18 | 9 | 24 | 24 | 1 |
| | | 56k | 30 | 13 | 116 | 41 | 3 |
| | | 92k | 17 | 9 | 425 | 44 | 1 |
| | 1k | 27k | 6 | 0 | 22 | 9 | 6 |
| | | 56k | 5 | 1 | 238 | 59 | 32 |
| | | 92k | 11 | 1 | 903 | 169 | 120 |
| | 10k | 27k | 0 | 0 | 4 | 0 | 0 |
| | | 56k | 0 | 0 | 360 | 77 | 172 |
| | | 92k | 10 | 0 | 379 | 116 | 508 |
