## Additional File 5 for "Genomic prediction with whole-genome sequence data in intensely selected pig lines"

### Genomic prediction accuracy by density of preselected variants in the prediction model

To test the impact of variant density on prediction accuracy, we preselected 10k, 25k, 75k, or 100k variants following the same criterion as described in the main text for Top40k. The difference of prediction accuracy between each set of preselected variants and Chip is shown, for all traits and lines (left) or by trait (right). Red dashed line at ‘no difference’. Regression coefficient (b) and p-value of training set size is provided, as well as the coefficient of determination ($R^2$) of the model. The linear model for the joint analyses included the trait effect.

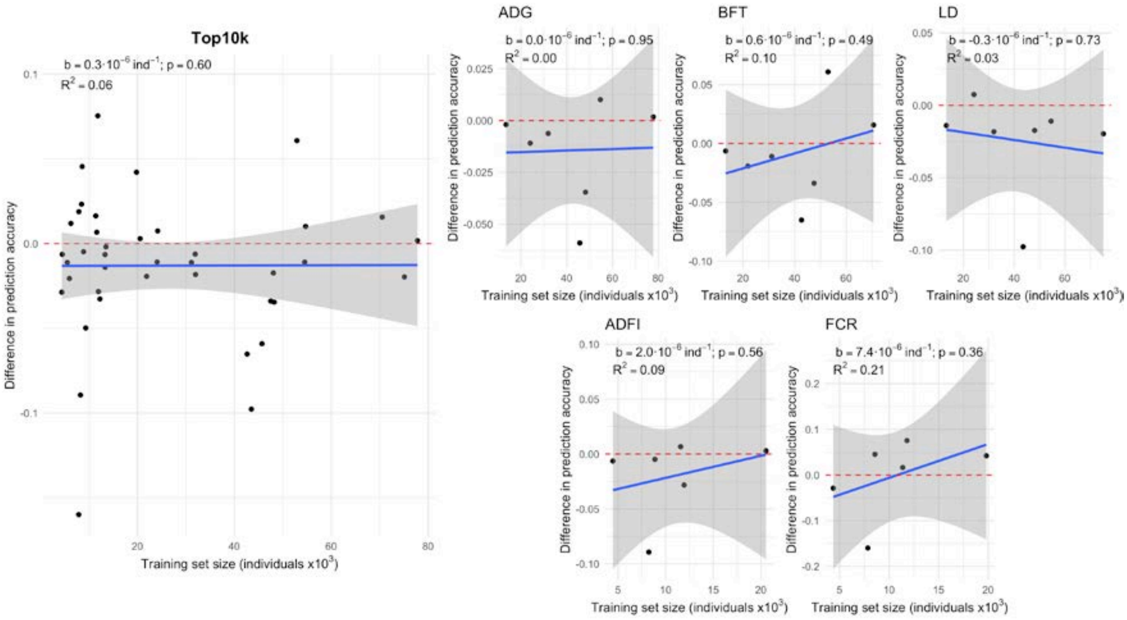


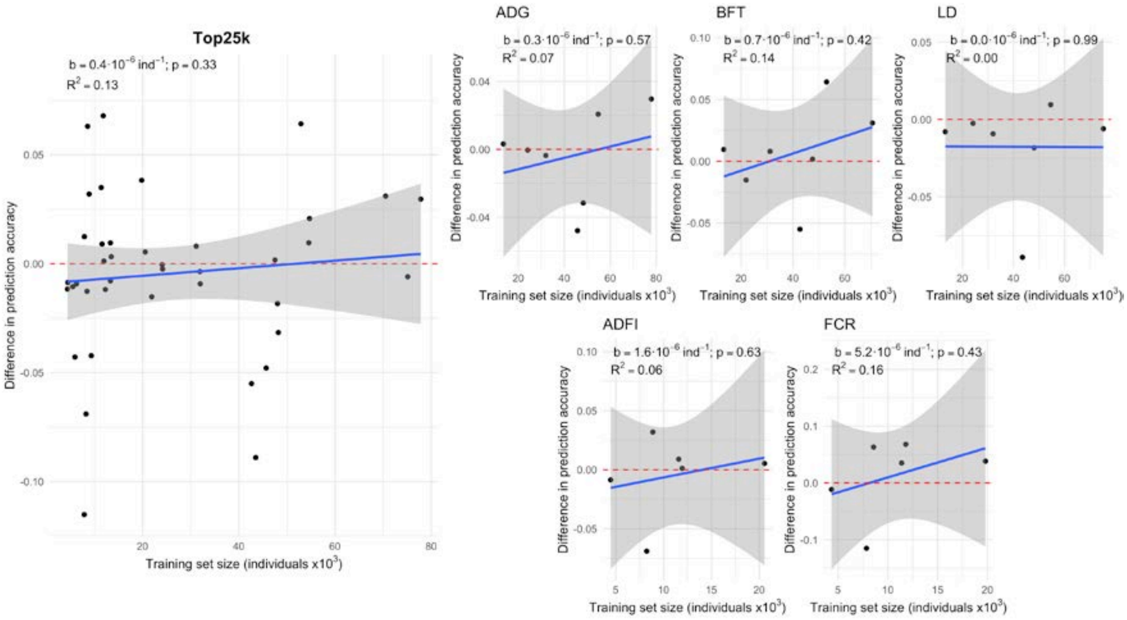

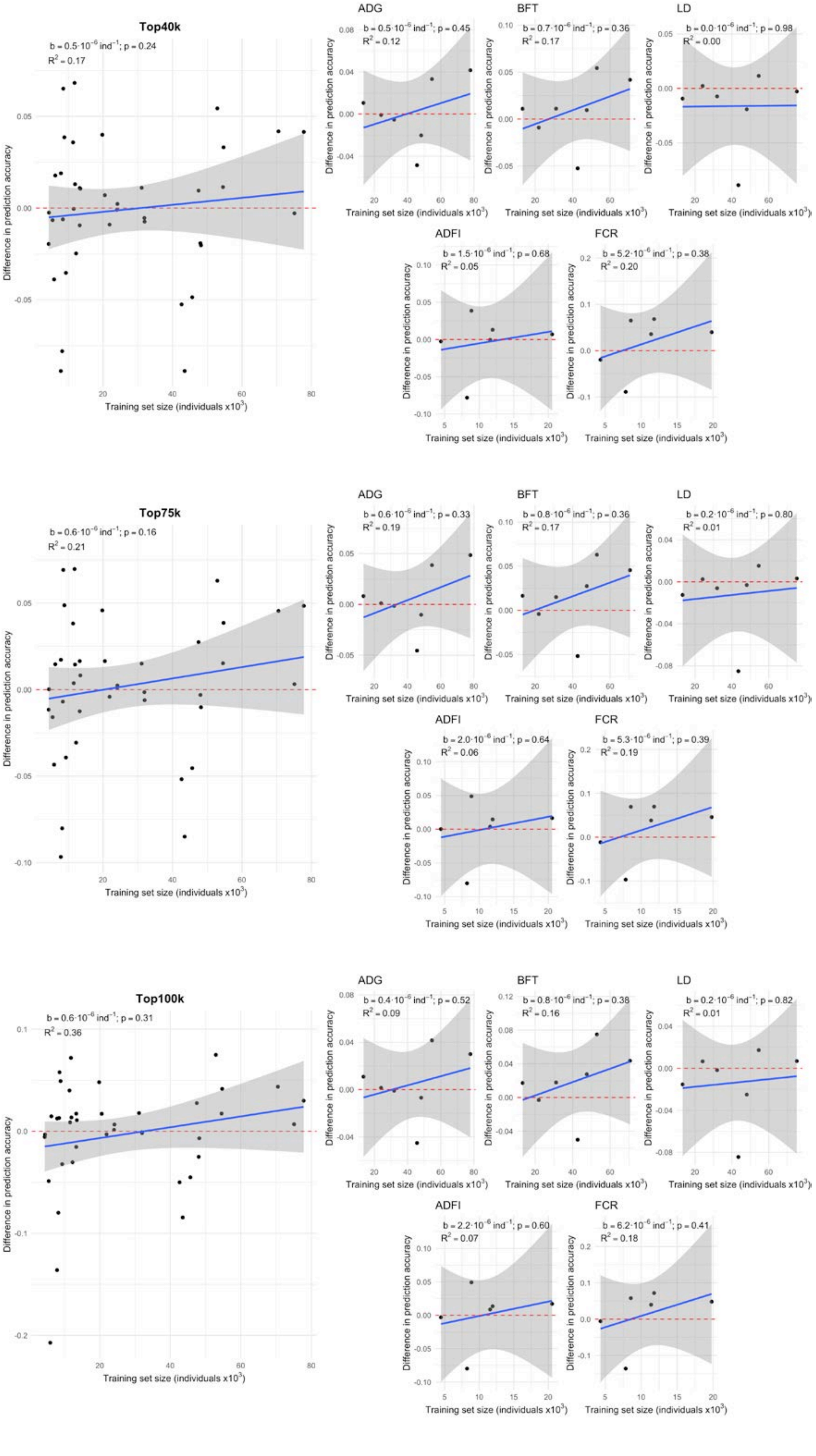
Top40k
b = 0.5·10^-6 ind^-1; p = 0.24
R^2 = 0.17
ADG
b = 0.5·10^-6 ind^-1; p = 0.45
R^2 = 0.12
BFT
b = 0.7·10^-6 ind^-1; p = 0.36
R^2 = 0.17
LD
b = 0.0·10^-6 ind^-1; p = 0.98
R^2 = 0.00
ADFI
b = 1.5·10^-6 ind^-1; p = 0.68
R^2 = 0.05
FCR
b = 5.2·10^-6 ind^-1; p = 0.38
R^2 = 0.20
Top75k
b = 0.6·10^-6 ind^-1; p = 0.16
R^2 = 0.21
ADG
b = 0.6·10^-6 ind^-1; p = 0.33
R^2 = 0.19
BFT
b = 0.8·10^-6 ind^-1; p = 0.36
R^2 = 0.17
LD
b = 0.2·10^-6 ind^-1; p = 0.80
R^2 = 0.01
ADFI
b = 2.0·10^-6 ind^-1; p = 0.64
R^2 = 0.06
FCR
b = 5.3·10^-6 ind^-1; p = 0.39
R^2 = 0.19
Top100k
b = 0.6·10^-6 ind^-1; p = 0.31
R^2 = 0.36
ADG
b = 0.4·10^-6 ind^-1; p = 0.52
R^2 = 0.09
BFT
b = 0.8·10^-6 ind^-1; p = 0.38
R^2 = 0.16
LD
b = 0.2·10^-6 ind^-1; p = 0.82
R^2 = 0.01
ADFI
b = 2.2·10^-6 ind^-1; p = 0.60
R^2 = 0.07
FCR
b = 6.2·10^-6 ind^-1; p = 0.41
R^2 = 0.18
Difference in prediction accuracy
Training set size (individuals x10^3)
