## Additional File 6 for "Genomic prediction with whole-genome sequence data in intensely selected pig lines"

### Genomic prediction accuracy of whole-genome sequence data in multi-line scenarios compared to the marker arrays in within-line scenarios

<u>ML-ChipPlusSign</u>

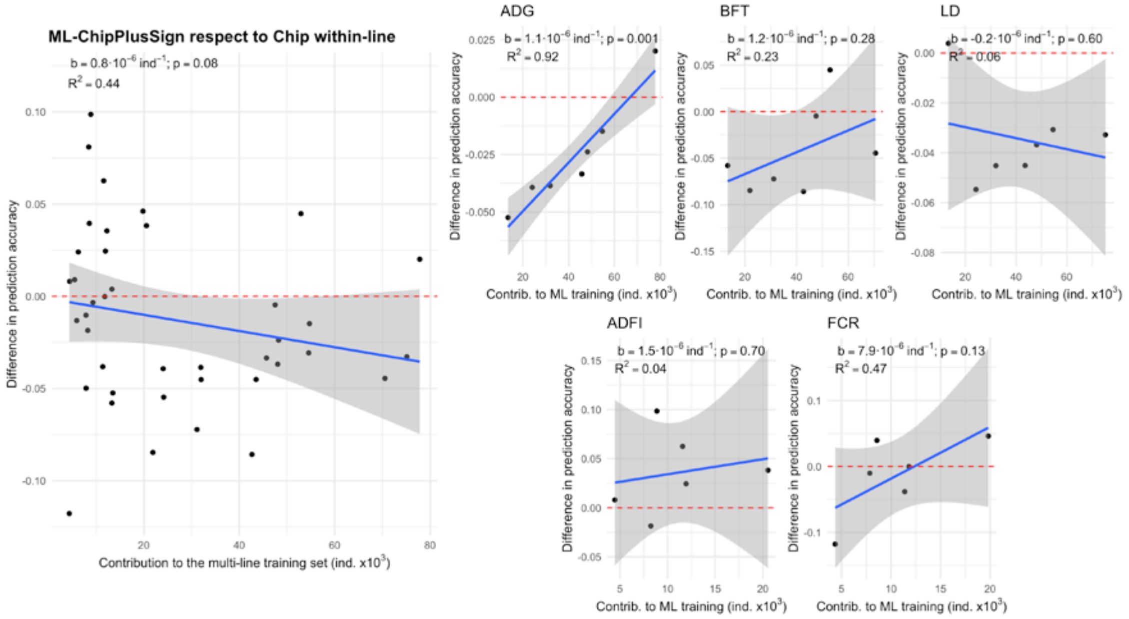


Difference of prediction accuracy between ML-ChipPlusSign and Chip in the within-line scenarios, for all traits and lines (left) or by trait (right). Red dashed line at ‘no difference’. Regression coefficient (b) and p-value of training set size is provided, as well as the coefficient of determination ($R^2$) of the model. The linear model for the joint analyses included the trait effect.

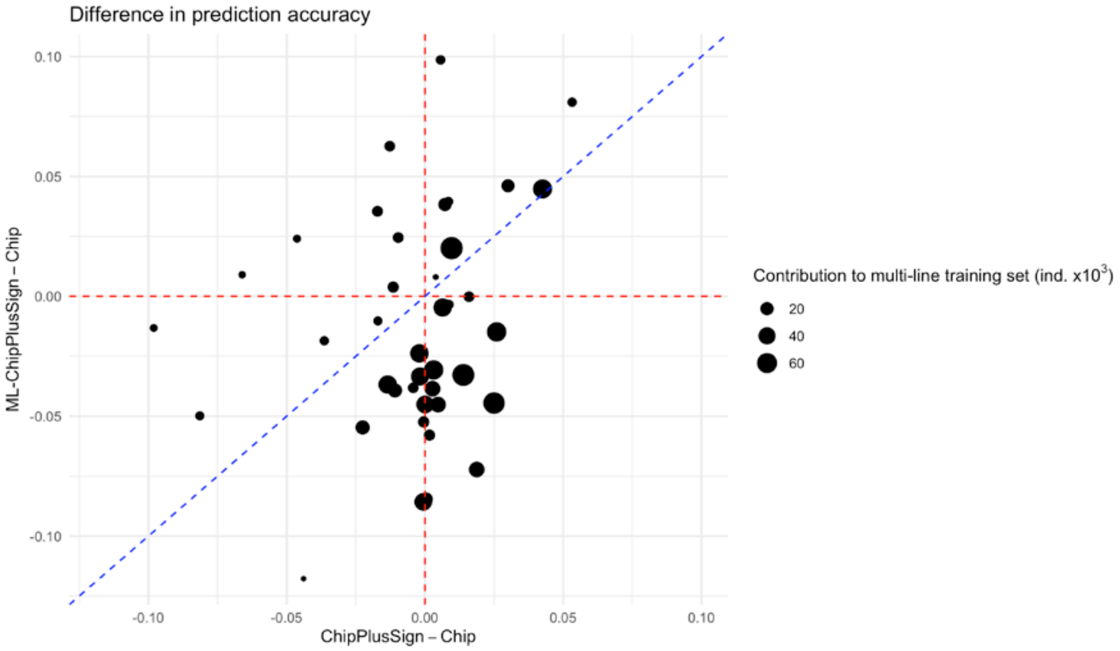


Comparison of the difference in genomic prediction accuracy in the multi-line scenarios (ML-ChipPlusSign) and in the within-line scenarios (ChipPlusSign), both respect to Chip in the within-line scenarios. Red dashed line at 1. Blue dashed line is the bisector.

ML-Top40k

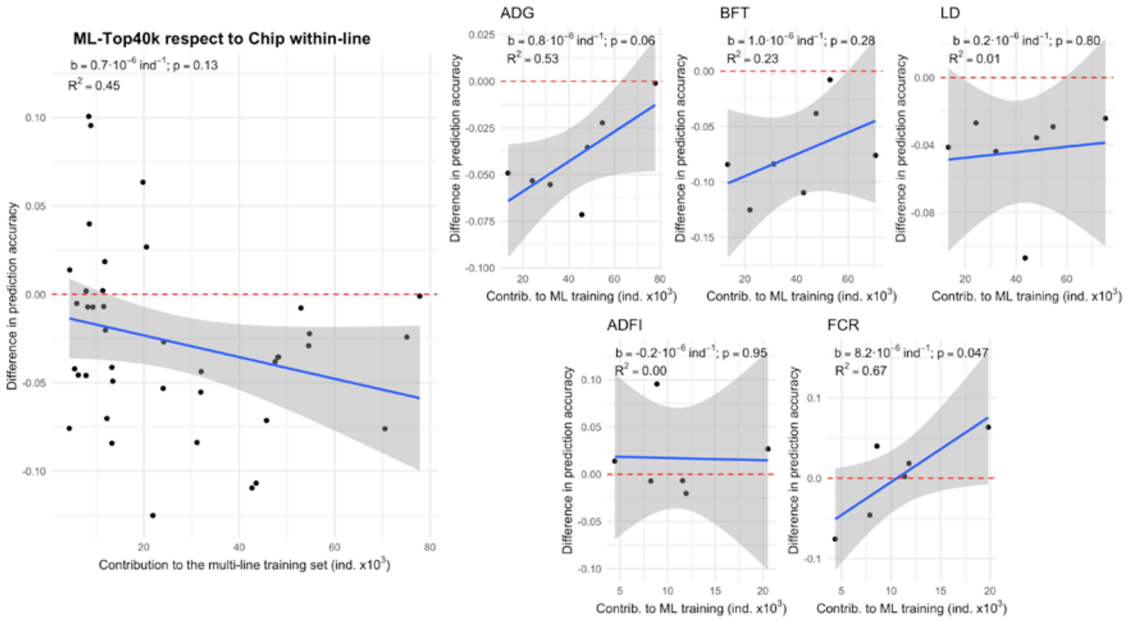


Difference of prediction accuracy between ML-Top40k and Chip in the within-line scenarios, for all traits and lines (left) or by trait (right). Red dashed line at ‘no difference’. Regression coefficient (b) and p-value of training set size is provided, as well as the coefficient of determination ($R^2$) of the model. The linear model for the joint analyses included the trait effect.

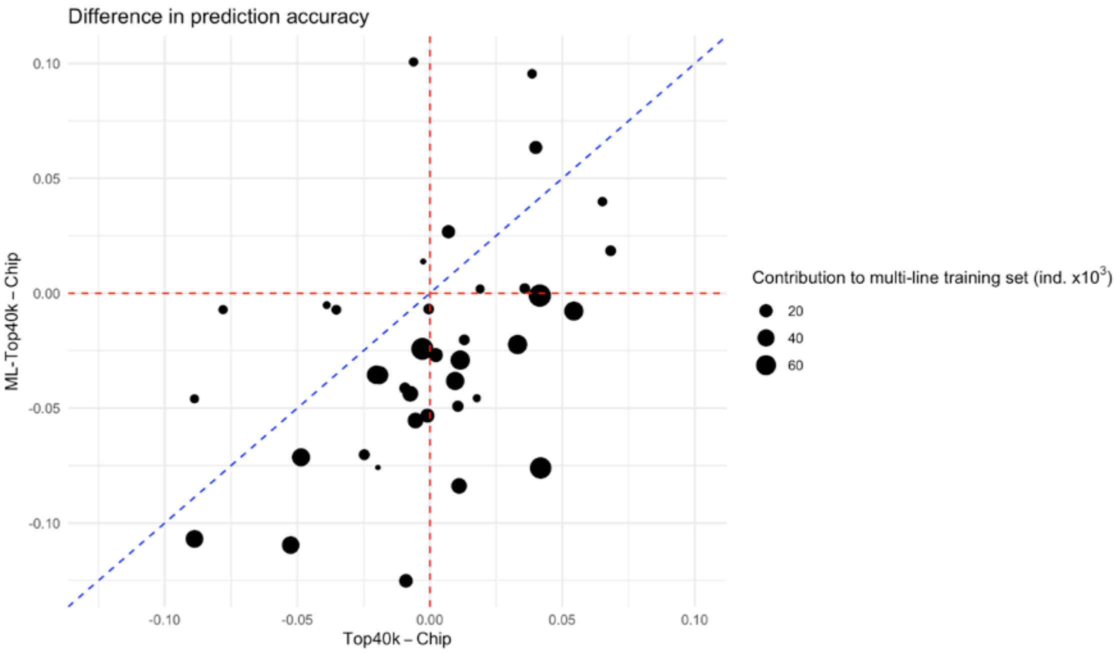


Comparison of the difference in genomic prediction accuracy in the multi-line scenarios (ML-Top40k) and in the within-line scenarios (Top40k), both respect to Chip in the within-line scenarios. Red dashed line at 1. Blue dashed line is the bisector.
