## Additional File 7 for "Genomic prediction with whole-genome sequence data in intensely selected pig lines"

**Additional File 6**

### Genomic prediction accuracy of ML-Top40k

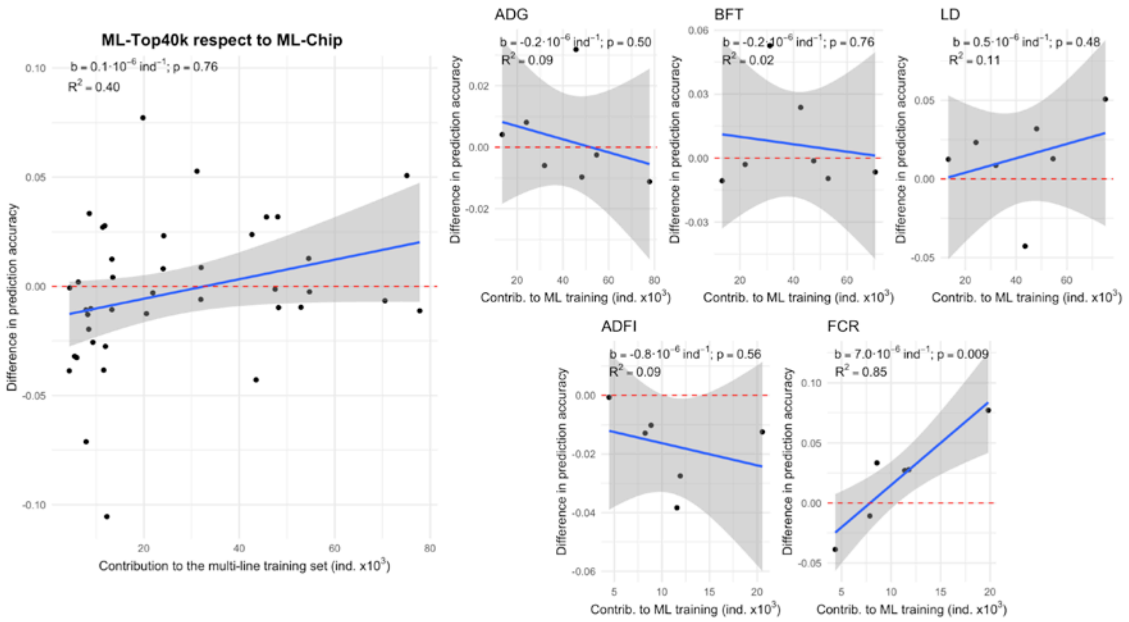


Difference of prediction accuracy between ML-Top40k and ML-Chip in the within-line scenarios, for all traits and lines (left) or by trait (right). Red dashed line at ‘no difference’. Regression coefficient (b) and p-value of training set size is provided, as well as the coefficient of determination ($R^2$) of the model. The linear model for the joint analyses included the trait effect.

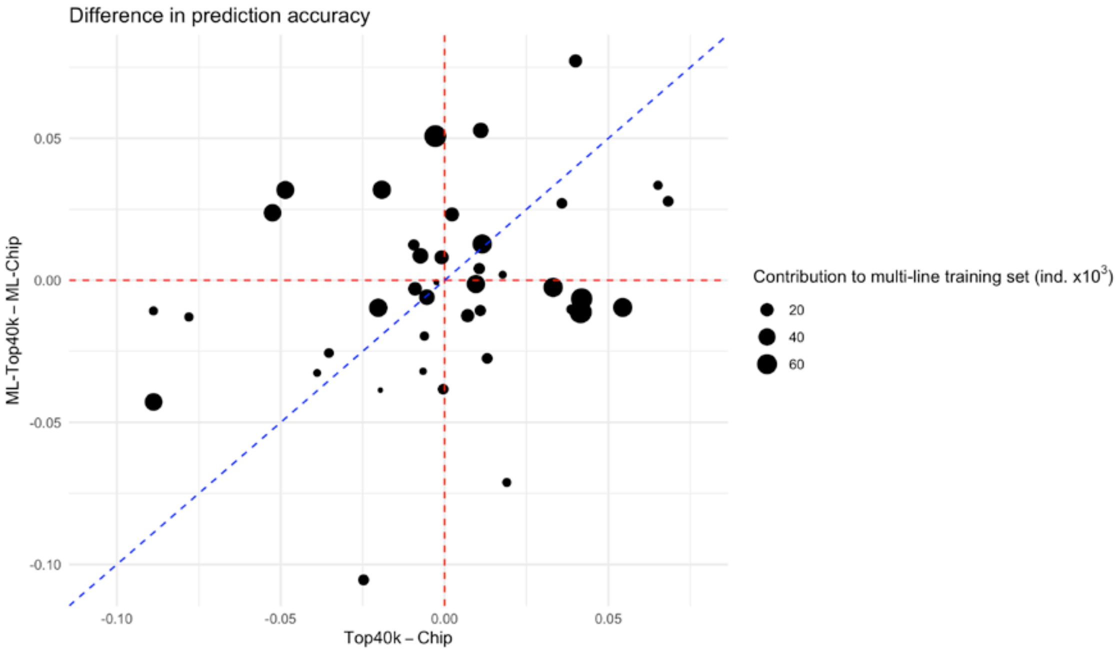


Comparison of the difference in genomic prediction accuracy in the multi-line scenarios (between ML-Top40k and ML-Chip) and in the within-line scenarios (between Top40k and Chip). Red dashed line at ‘no difference’. Blue dashed line is the bisector.
