## Additional File 8 for "Genomic prediction with whole-genome sequence data in intensely selected pig lines"

### Genome-wide association study on backfat thickness with marker array or whole-genome sequence data

#### Methods

To assess whether variants from the whole-genome sequence data (WGS) could provide a finer mapping of causal variants than marker array data (Chip), we performed a genome-wide association study (GWAS) for backfat thickness (BFT) in line A using either the marker array data or the WGS. The GWAS was performed as described in the main text. Due to computational limitations, the genomic relationship matrix **K** was calculated using only imputed genotypes in the marker array regardless of whether the association study involves the variants in the marker array or WGS. We used a p-value threshold of $p \leq 10^{-6}$ for associations of marker array data, based on Bonferroni's multiple test correction assuming that the markers from the marker array were independent.

#### Results

To illustrate the performance of GWAS using WGS compared to marker arrays, we examined the GWAS results for chromosome 1 (as an example) and for six genomic regions with significant associations. The main genomic regions and candidate genes associated to BFT detected with marker arrays in the same genetic lines were reported by Gozalo-Marcilla et al. (2021; doi: 10.1186/s12711-021-00671-w). As a reference, candidate functional genes for each region are indicated according to Gozalo-Marcilla et al. (2021).

Using marker array, we identified 6 genomic regions with significant associations. Using WGS, we confirmed 3 of these genomic regions that co-located to candidate genes *MC4R*, *DOLK*, and *DGKI* or *PTN*. The most associated variants in each of these genomic regions were located outside the coding region of these putative causal genes. These signals sometimes had very strong evidence of association for some variants that were relatively distant from our candidate functional gene, which could cast doubts about the fine-mapping of the causal mutation. The region at SSC18, 9–13 Mb, contained two candidate genes *DGKI* and *PTN*, but the WGS revealed significantly associated variants within *DGKI* and none within *PTN*, despite that the strongest associations were away from both genes at 10.5-11 Mb. Using the WGS we also detected 24 additional genomic regions that contained candidate genes such as *CYB5R4*, *IGF2*, and *LEPR*. These genes were previously detected in other lines using marker array data but not in this one (Gozalo-Marcilla et al., 2021), sometimes because there were no markers for the associated region in our marker array data (SSC2, 0–4 Mb). The region at SSC1, 52.5–53.5 Mb, showed many significant variants that encompassed not only the previously identified candidate gene *CYB5R4*, but also *MRAP2* (annotated with functions on feeding behavior and energy homeostasis). In contrast, candidate gene *LEPR* was located within the region at SSC6, 146.5–147.0 Mb, where many significant variants were located, although the most significant variants were not in the coding regions of the gene. For many of the other genomic regions, it was difficult to pinpoint a candidate gene with the available information or there were no annotated genes.

#### Figures

In red, results for the variants in the marker array (Chip); in black, results for the WGS. The blue dashed line indicates $p=10^{-6}$. Below each plot of a genomic region, annotated genes from Ensembl 104.

<u>Chromosome SSC1</u>

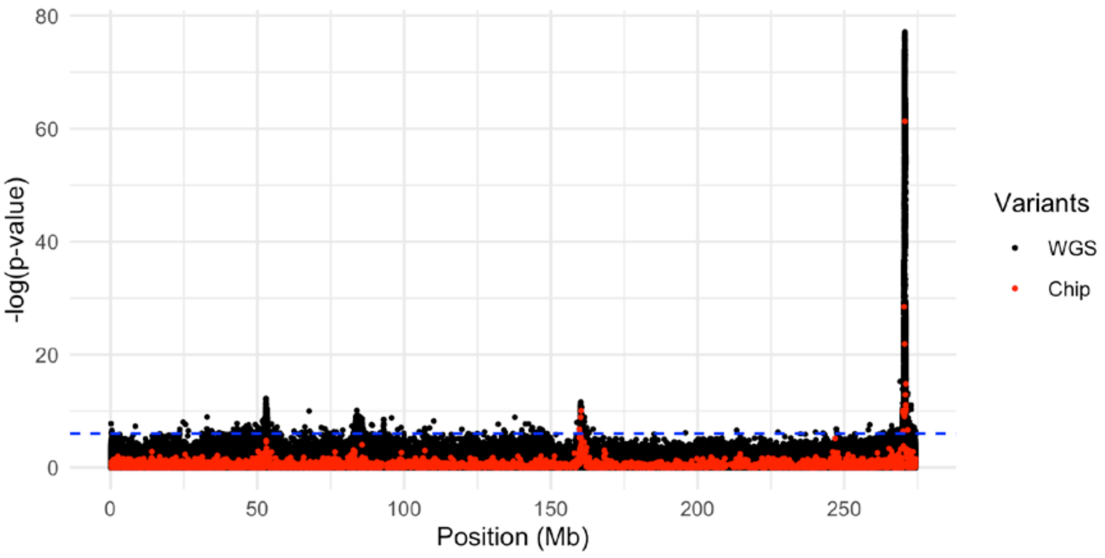


<u>SSC1, 51–55 Mb. Candidate functional genes: *CYB5R4*</u>

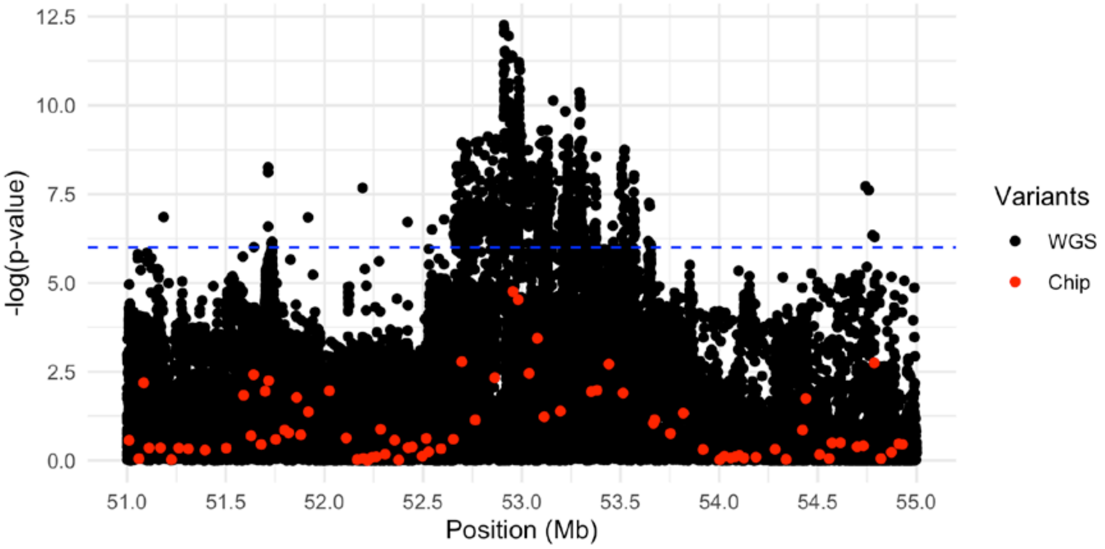


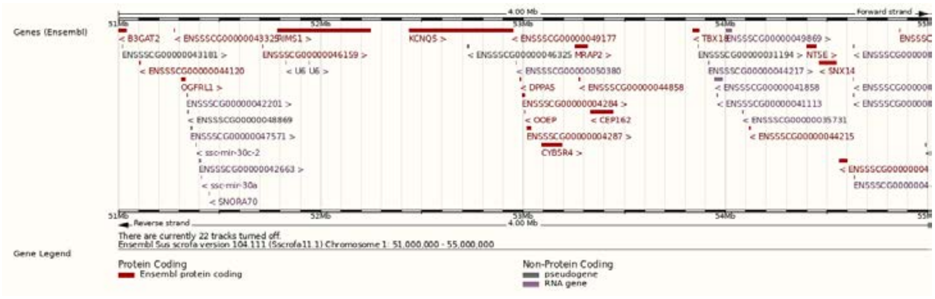

SSC1, 158–162 Mb. Candidate functional gene: *MC4R*

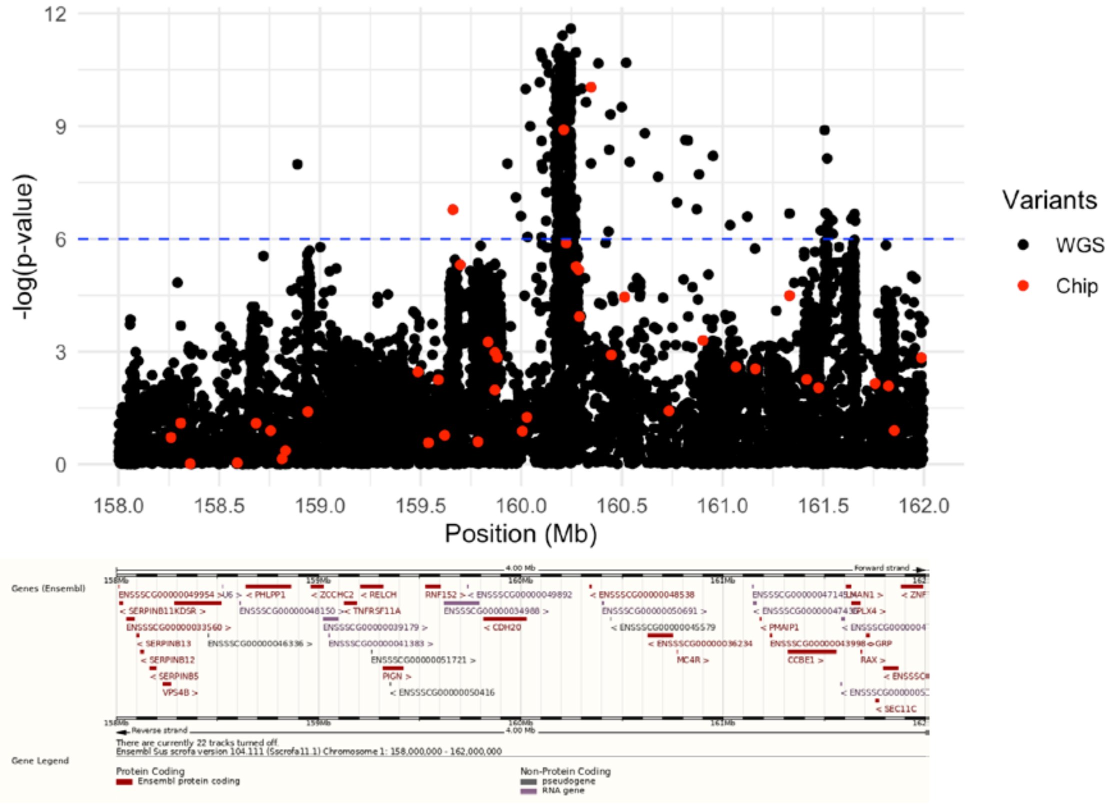


SSC1, 268–272 Mb. Candidate functional gene: *DOLK*

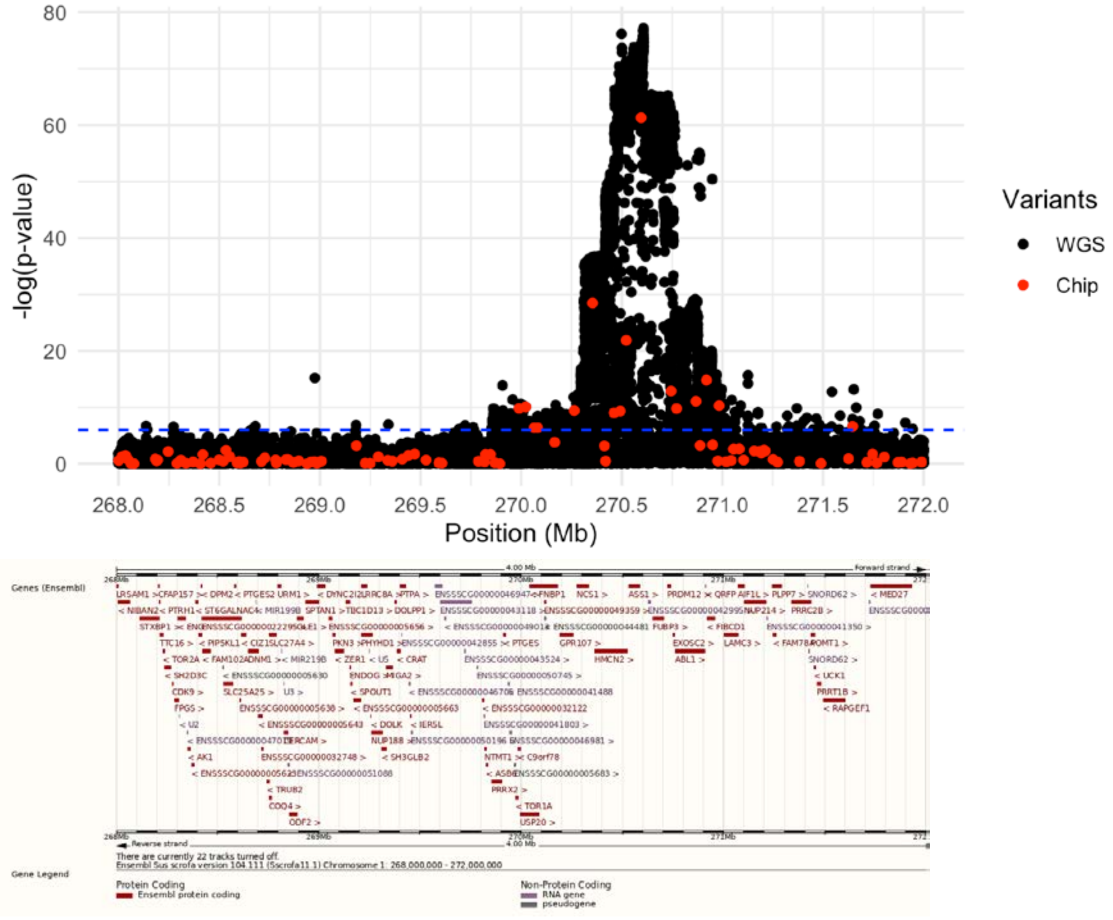

<u>SSC2, 0–4 Mb. Candidate functional gene: *IGF2*</u>

*IGF2* not annotated in Ensembl 104. Position from NCBI: 1.47-1.50 Mb

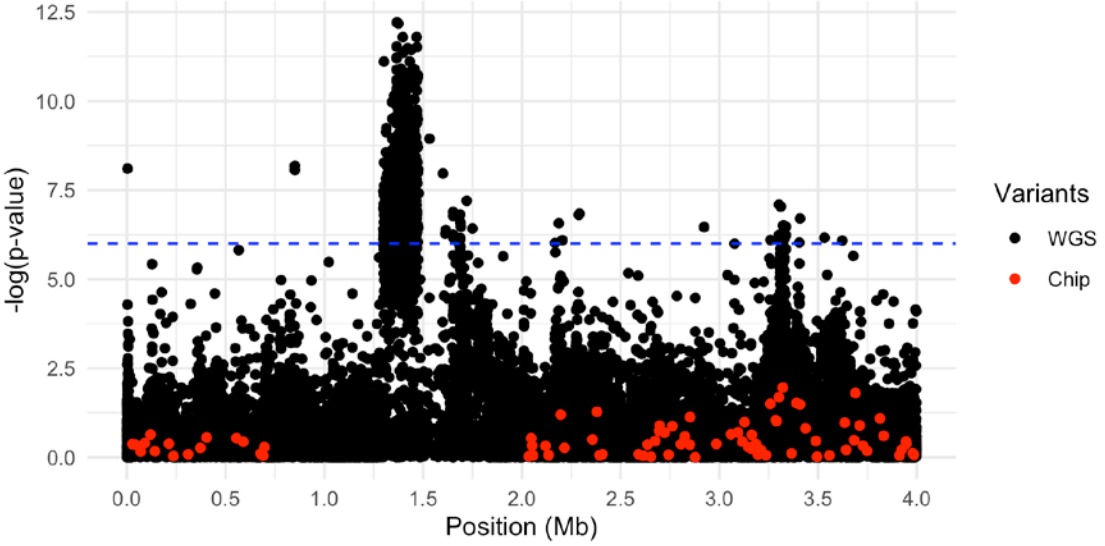


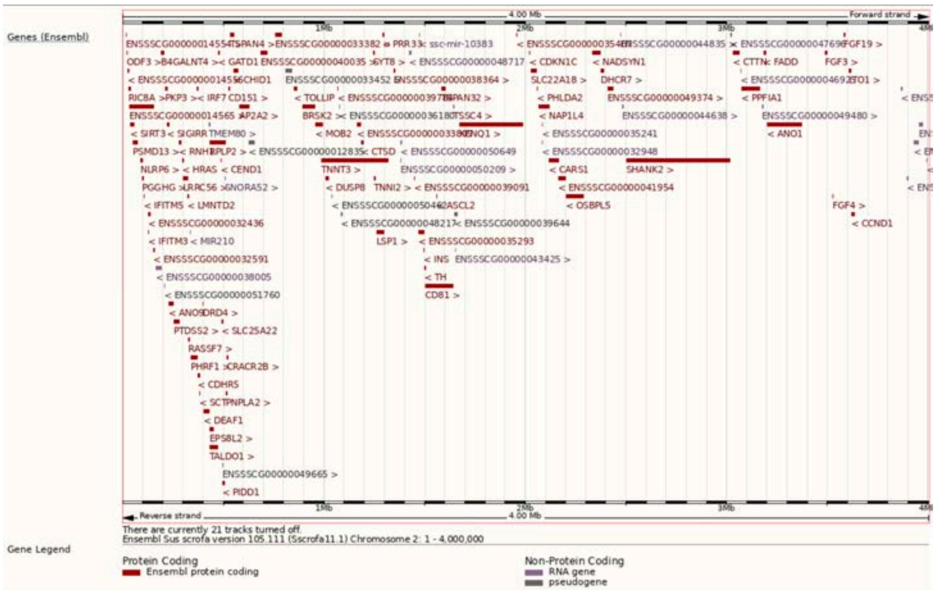

SSC6, 145–149 Mb. Candidate functional gene: *LEPR*

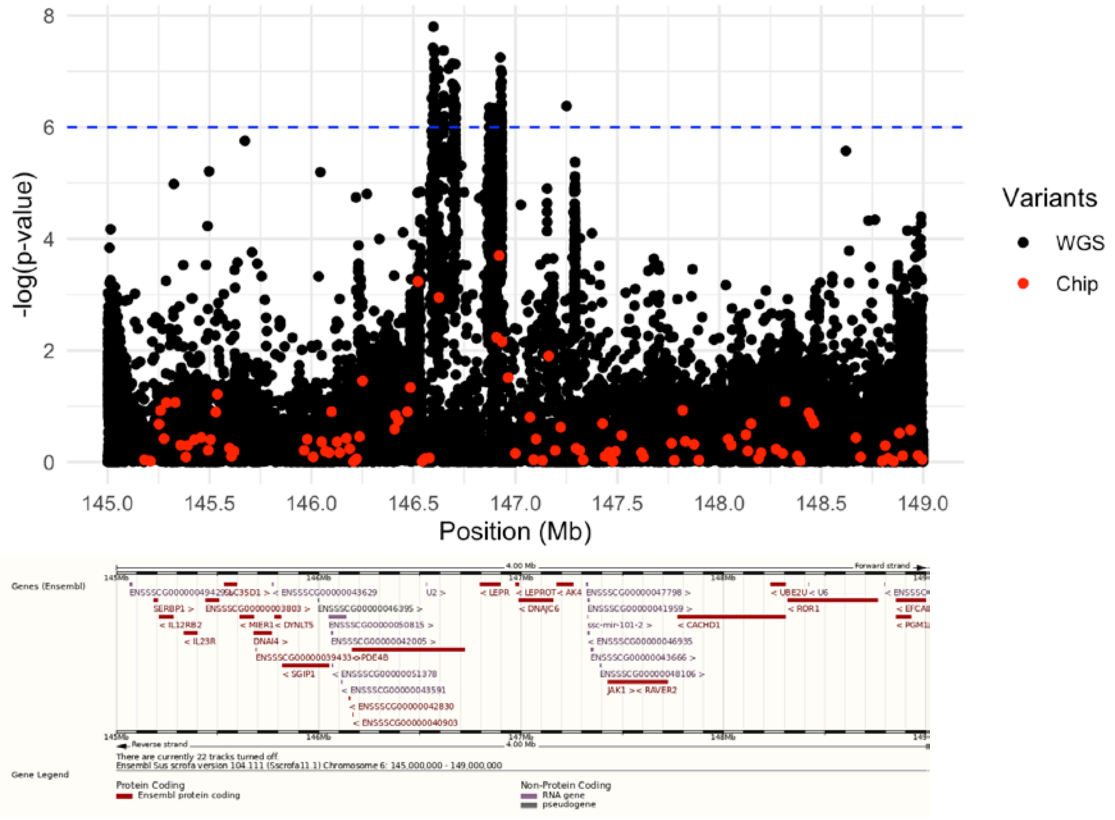


SSC18, 9–13 Mb. Candidate functional gene: *DGKI* and *PTN*

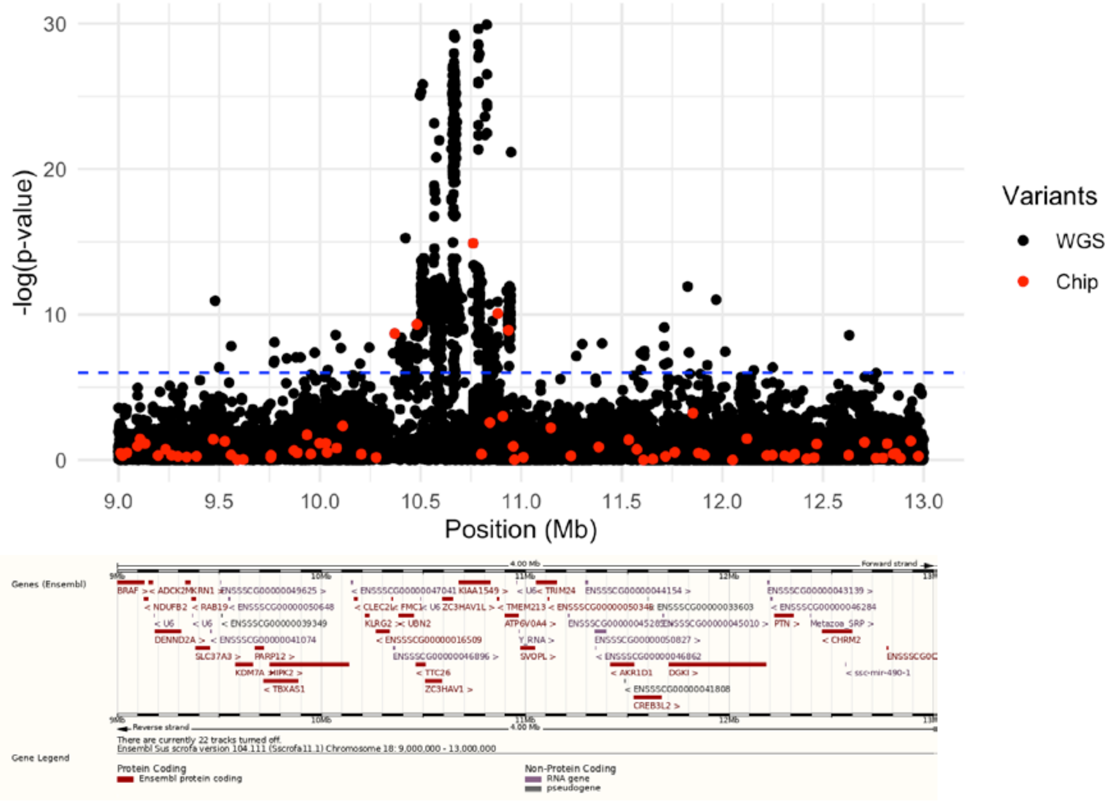
