## Additional File 9 for "Genomic prediction with whole-genome sequence data in intensely selected pig lines"

### Genomic prediction accuracy and bias when preselection of variants and training of the predictive equation are performed in two mutually exclusive subsets

#### Methods

We reanalysed the data after splitting the training set into two mutually exclusive subsets, one for GWAS to preselect the predictor variants and one for training the predictive equation. The GWAS subset was defined by randomly selecting either 10% or 50% of the individuals in the original training set. Those individuals were excluded from the subset used for training the predictive equation afterwards. Thus, three cases are considered: (i) 10% for GWAS-based preselection of variants + 90% for training of the predictive equation; (ii) 50% for GWAS-based preselection of variants + 50% for training of the predictive equation; and (iii) all individuals for both GWAS-based preselection of variants and training of the predictive equation. The rest of analyses were performed as described in the main text.

#### Results

Splitting the original training set into two mutually exclusive subsets, one for the GWAS-based preselection of the variants and one for the training of the predictive equation, did not improve the performance of the WGS for genomic prediction compared to using the same set for both preselecting variants and training the predictive equation. For ChipPlusSign, this strategy reduced the bias but the difference between prediction accuracy of WGS and the marker array decreased too, probably because of the smaller subset available for training the predictive equation.

#### Figures

Left: Difference of prediction accuracy between each set of predictor variants and Chip. Red dashed line at ‘no difference’. Regression coefficient (b) and p-value of training set size is provided, as well as the coefficient of determination ($R^2$) of the model. The linear model included the trait effect. Right: Comparison of the bias between each set of predictor variants and Chip. Red dashed lines at the ideal value of 1. Blue dashed line is the bisector.

ChipPlusSign

(i)

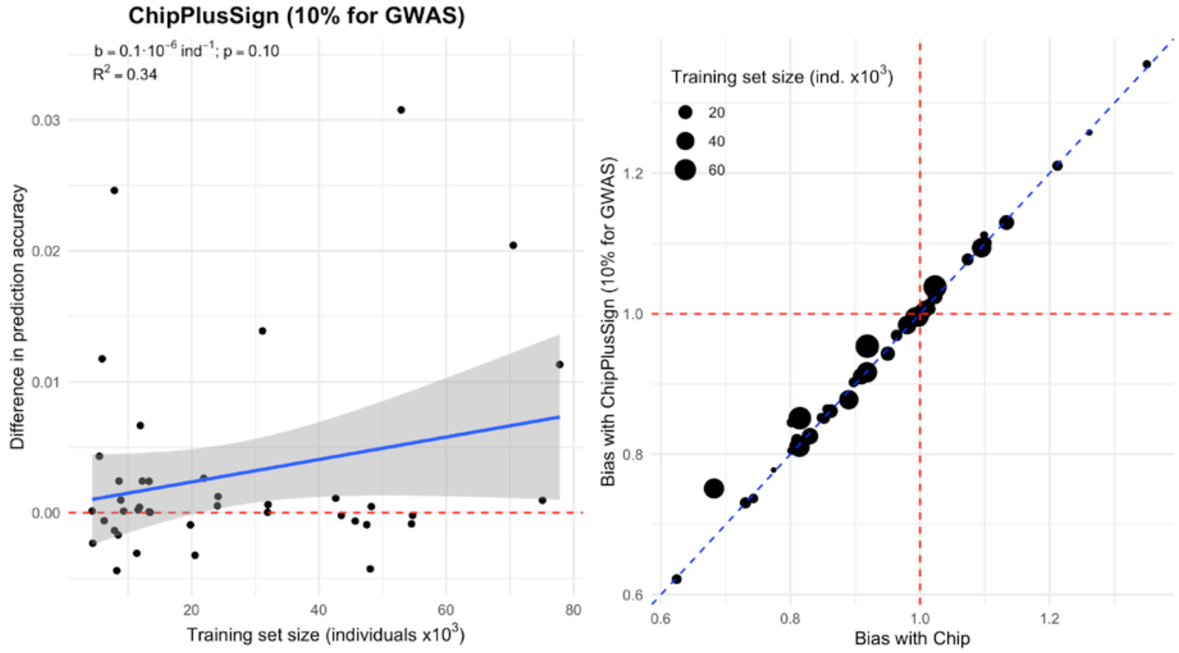


(ii)

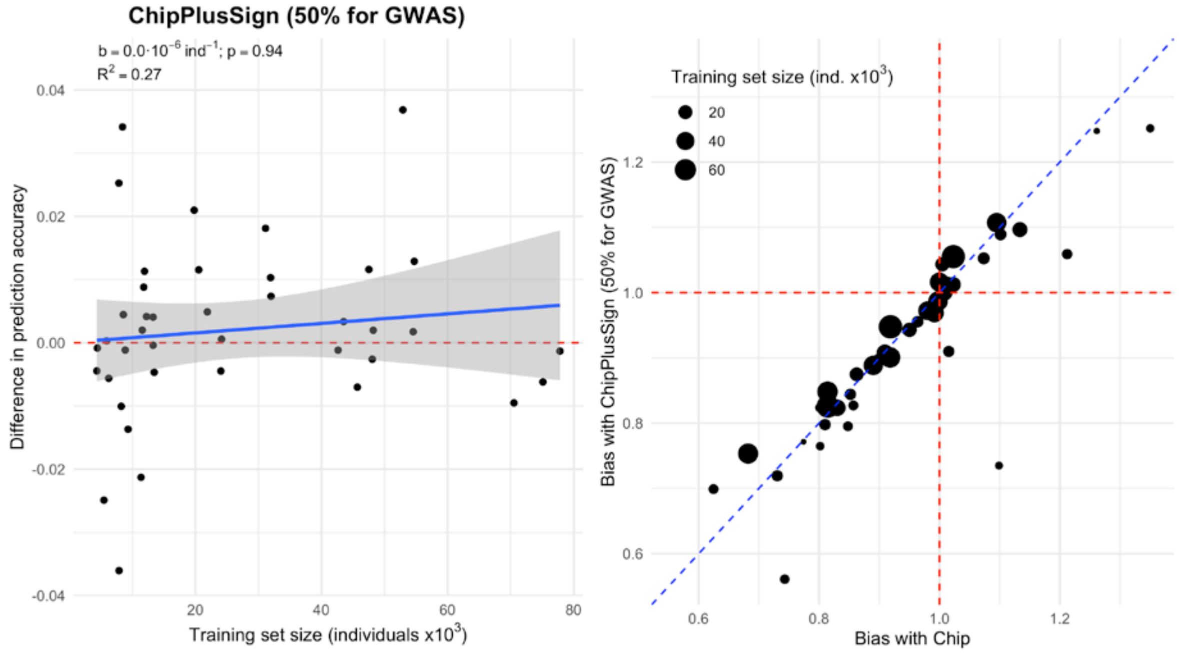

(iii)

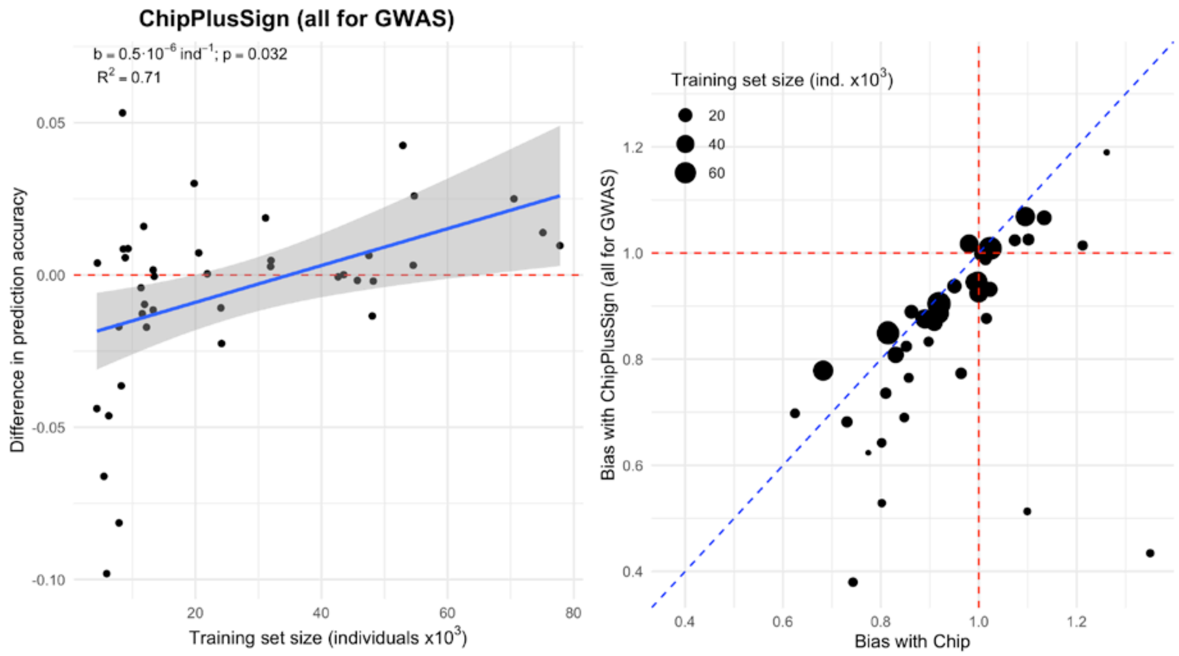


<u>Top40k</u>

(i)

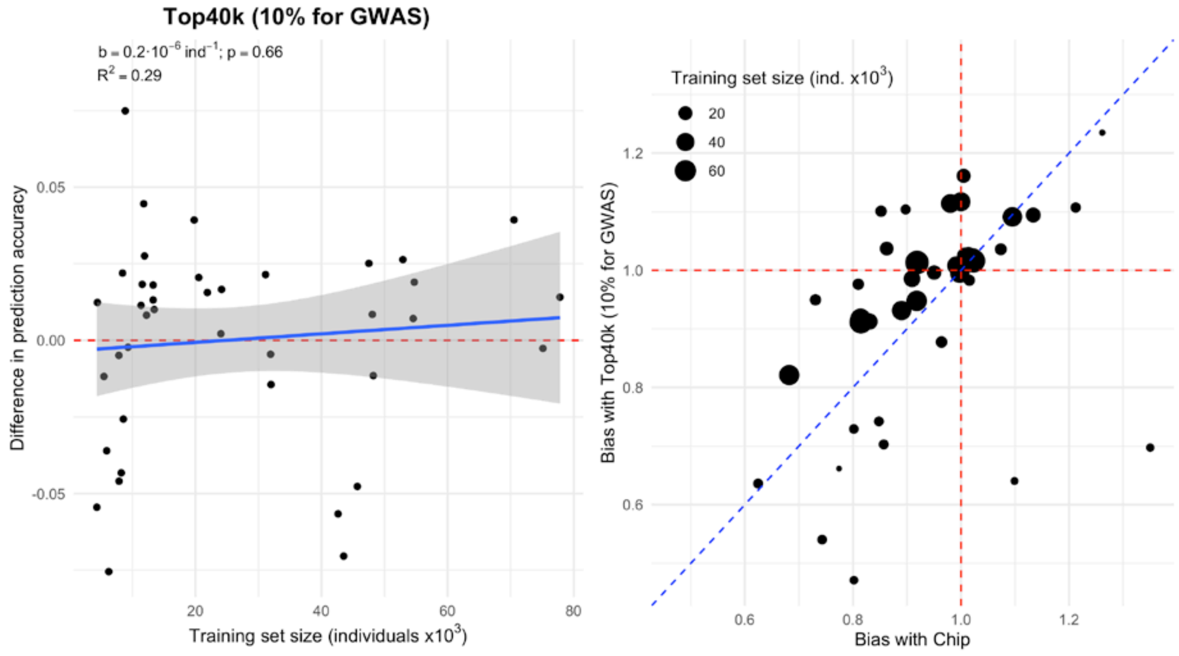

(ii)

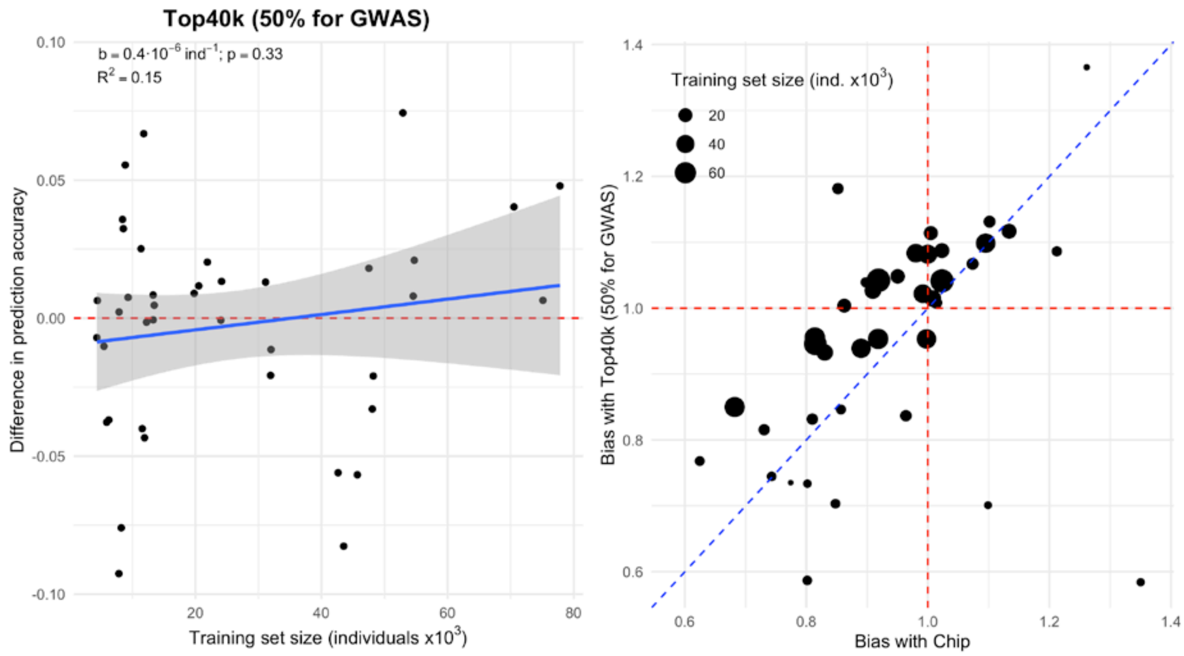
Top40k (50% for GWAS)
b = 0.4·10$^{-6}$ ind$^{-1}$; p = 0.33
$R^2$ = 0.15
Difference in prediction accuracy
Training set size (individuals x10$^3$)
0.10
0.05
0.00
-0.05
-0.10
20
40
60
80
Training set size (ind. x10$^3$)
20
40
60
Bias with Top40k (50% for GWAS)
Bias with Chip
1.4
1.2
1.0
0.8
0.6
0.6
0.8
1.0
1.2
1.4

(iii)

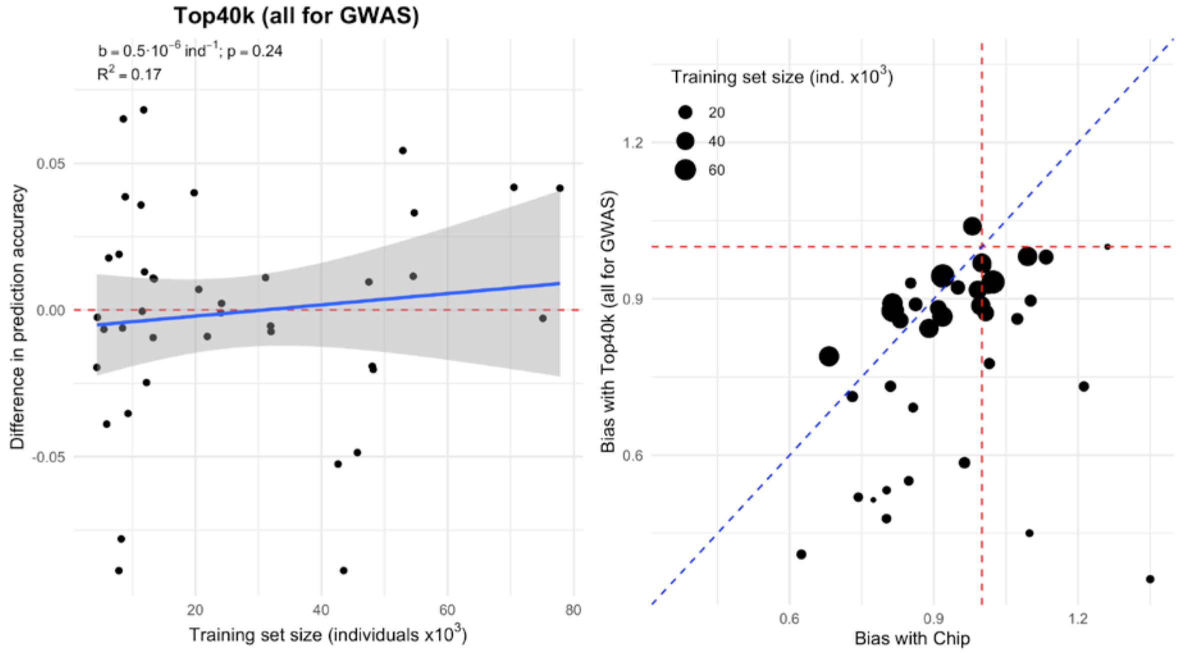
Top40k (all for GWAS)
b = 0.5·10$^{-6}$ ind$^{-1}$; p = 0.24
$R^2$ = 0.17
Difference in prediction accuracy
Training set size (individuals x10$^3$)
0.05
0.00
-0.05
20
40
60
80
Training set size (ind. x10$^3$)
20
40
60
Bias with Top40k (all for GWAS)
Bias with Chip
1.2
0.9
0.6
0.6
0.9
1.2
